# Fungal infection alters collective nutritional intake of ant colonies

**DOI:** 10.1101/2023.10.26.564092

**Authors:** Enikő Csata, Alfonso Pérez-Escudero, Emmanuel Laury, Hanna Leitner, Gérard Latil, Jürgen Heinze, Stephen J. Simpson, Sylvia Cremer, Audrey Dussutour

## Abstract

In many animals, parasitic infections impose significant fitness costs [1–6]. Animals are known to alter their feeding behavior when infected to help combat various parasites [7–12]. For instance, they can adjust nutrient intake to support their immune system [13,14]. However, parasites can also manipulate host foraging behavior to increase their own development, survival and transmission [15–18]. The mechanisms by which nutrition influences host-parasite interactions are still not well understood. Until now, studies that examine the impact of diet on infection have mainly focused on the host, and less on the parasite [12,13, 19–25]. Using Nutritional Geometry [26], we investigated the role of key nutrients: amino acids and carbohydrates, in a host-parasite system: the Argentine ant, *Linepithema humile,* and the entomopathogenic fungus, *Metarhizium brunneum*. We first established that the fungus grew and reproduced better on diets comprising four times less amino acids than carbohydrates (1:4 AA:C ratio). Second, when facing food combinations, the fungus exploited the two complementary food resources to reach the same performance as on this optimal diet, revealing the ability of fungal pathogens to solve complex nutritional challenges. Third, when ants were fed on this optimal fungal diet, their lifespan decreased when healthy, yet not when *Metarhizium*-infected, compared to their favored carbohydrate-rich diet. Interestingly, when the ants were given a binary choice between different diets, the foragers of uninfected colonies avoided intake of the fungal optimum diet, whilst choosing it when infected. Experimental disentanglement of full pathogenic infection and pure immune response to fungal cell wall material, combined with immune measurements, allowed us to conclude that this change of nutritional choice in infected ants did not result from pathogen manipulation but likely represents a compensation of the host to counterbalance the cost of using amino acids during the immune response. The observed change in foraging behavior in infected colonies towards an otherwise harmful diet (self-medication), suggests a collective compensatory mechanism for the individual cost of immunity. In short, we demonstrated that infected ants converge on a diet that is proven to be costly for survival in the long term but that could help them fight infection in the short term.

**Highlights:**

1. The insect-pathogenic fungus *Metarhizium brunneum* performs best on protein-rich diets and is able to solve complex nutritional challenges
2. While harmful to healthy ants, protein-rich diets did not shorten infected ants’ lifespan
3. Contrary to healthy ants, when given a choice, infected and immune-stimulated ants choose a protein-rich diet

## RESULTS

We used Nutritional Geometry (NG, [26]) to study how an entomopathogenic fungus affects the nutritional choices of ant colonies. First, we defined independently the intake target (optimal diet) of the pathogen, the entomopathogenic fungus *Metarhizium brunneum*, on agar-based artificial diets, and of uninfected and infected ant hosts, *Linepithema humile*. Second, we challenged the fungus with various food pairings to test whether the fungus is able to exploit two complementary food resources to maximize its performance. Next, we studied how infected and uninfected hosts performed when confined to diets with differing amounts of amino acids and carbohydrates. Then, we offered both infected colonies and uninfected colonies a choice between an amino acids-rich and a carbohydrate-rich food, to investigate whether foragers adapt their foraging strategy in response to the infection. A change in colony nutrient intake may indicate either parasite manipulation of host foraging strategies to promote their own growth, or a host strategy to promote its immune system to better fight infection.

To disentangle these two hypotheses, we injected ant workers with β-1,3 glucan from cell walls (zymosan), known elicitors of the ant immune system [27,28]. By determination of immune gene expression, we confirmed that this injection activated antimicrobial peptide production in Argentine ants. We then offered these immune-stimulated, as well as *Metarhizium*-infected ants, a choice between two foods with differing amounts of amino acids and carbohydrates and compared their food intake responses.

### The fungal pathogen *Metarhizium brunneum* thrives on a high-protein diet

We first established the optimal diet composition of the fungal pathogen *Metarhizium brunneum* for the two major macronutrients, essential amino acids (AA), the building blocks of proteins, and carbohydrates (C). To this end, we confined the fungus to one of 18 agar-based artificial diets, varying in both their AA:C ratio and the total concentration of AA+C (*see* Table S1 for AA:C ratios and Table S2 for AA+C for total concentrations). For each diet, we then measured the extent of fungal expansion (growth rate: area mm^2^) after 14 and 30 days, when fungal conidiospores (hereafter abbreviated as spores) were harvested and counted.

We found that the number of spores produced per unit of surface was highest on the diet with an AA:C ratio of 1:4, i.e. comprising four times more carbohydrates than essential amino acids (Fig. 1*A*; Fig. S1; Table S3; p < 0.001). The fungus also grew to cover the largest area on this 1:4 diet (Fig. 1*B*,*C* and Fig. 1*D*), which was also the case after 14 days (Fig. S2; Table S4; p < 0.001) and 30 days (Fig. S3; Table S5; p < 0.001). Together, this meant that fungal reproduction was highest on a diet comprising four times more carbohydrates than essential amino acids (Fig. 1*E*; Table S6).

**Figure 1.**
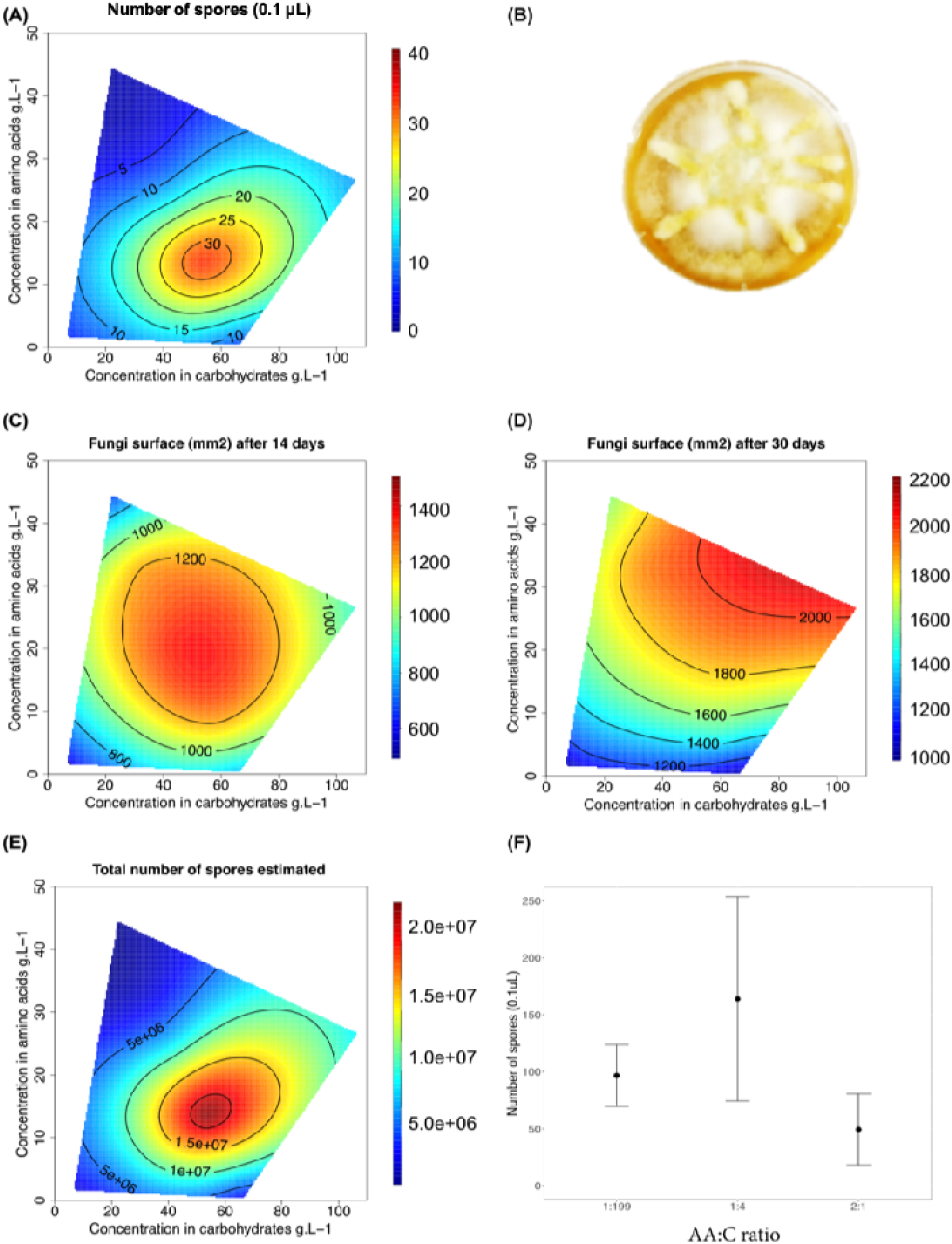
Fungal growth and spore production on different nutrient compositions (artificial diets, Table S1). **Figure A, C-E.** Performance responses. **A.** The number of spores produced by *M. brunneum* per µL (volume = 0.1µL) was quantified for the fungus confined for 30 days to one out of 18 diets varying in both the ratio and total amount of amino acids and carbohydrates. The response surface regression analyses yielded a significant relationship: p < 0.001. **B.** Photograph of fungal growth on a 1:4 artificial diet after 30 days. **C and D**. Fungal growth (mm^2^) was determined for fungus grown for 14 and 30 days on one out of 18 diets. The response surface regression analyses showed significant relationships for day 14 (p < 0.001) and day 30 (p < 0.001). The number of replicates per diet is between 11 and 20. **E.** The total number of spores calculated for fungi confined for 30 days to one of 18 diets. For each replicate, the number of spores harvested by the surface unit was multiplied by the total surface covered by the fungus. The response surface regression analyses yielded a significant relationship (p < 0.001). **F.** Effect of past diet on spore production. The mean number of secondary spores (spores that were produced when reared under previous fungal diet restrictions, now all plated on standard medium) harvested (per 0.1µL) on a standard medium as a function of the diet (AA:C ratio, 1:4, 1:199 and 2:1) used to produce the primary spores (N = 5 to 6 replicates per diet). Linear model F_2,13_ = 4.57, p = 0.031 followed by all-pairwise post-hoc comparisons (Tukey method; 1:199 vs 1:4 p = 0.696, 1:199 vs 2:1 p = 0.139, 1:4 vs 2:1 p = 0.027). Error bars indicate the 95%CI.

For nutrient concentration, we found that the fungus produced more spores on diets with intermediate concentrations of AA+C than on very low- or high-concentrated ones (Fig. 1*A*; Fig. S4; Table S3; p < 0.001). Similarly, fungal expansion was greatest on intermediate-concentrated diets on day 14 (Fig. S5; Table S4; p < 0.001). On day 30, the fungus had expanded across a greater area on both, the intermediate and high-concentrated diets. Therefore, we found no indication that the fungus would increase the surface in contact with the food to compensate for low nutrient availability, but, on the contrary, could expand more with higher nutrient amounts available (Fig. 1*C*; Fig. S6; Table S5; p < 0.001).

Since not only spore quantity but also quality will affect pathogen fitness, we next measured if viability and germination ability differed between spores that had been produced under different nutrient regimes, but were then all grown on the same standard medium (Malt Extract Agar). For this second experiment, we used spores originating from the two most extreme diets (high carbohydrate diet 1:199 and high essential amino acid diet 2:1), as well as from the fungal optimal diet maximizing spore production (1:4). We found that the number of spores growing per unit of surface was significantly higher for the spores originating from the optimal 1:4 diet ratio compared to spores originating from the high amino acid-biased diet (2:1) diet (Fig. 1F; Table S7; 1:4 vs 2:1 p = 0.02). Spores originating from the high carbohydrate-biased diet (1:199) showed intermediate growth (1:199 vs 1:4 p = 0.69, 1:199 vs 2:1 p = 0.13).

We further tested if the fungal pathogen could combine nutrients of two highly imbalanced but complementary nutrient sources, by plating fungus on six pairings of nutrient sources differing in their AA:C ratio (either 1:199 vs. 2:1, 1:49 vs. 2.1, 1:24 vs. 2:1, 1:16 vs. 2:1, 1:12 vs. 2:1, or 1:8 vs. 2:1, *see* schematic design Fig. S7; Experiment 3). After 30 days, the fungus was able to produce the same number of spores per unit of surface in each diet pairing as on the 1:4 diet in the no-choice experiment (Fig. 2*A*; Fig. S7; Table S8; p = 0.15). At this time, the overall area covered by the fungus was also the same as on the 1:4 diet in the no-choice experiment (Fig. 2*C*), yet it was differently distributed over the two diet types: the area on the high amino acid-biased diet (2:1) was always smaller than that on its paired carbohydrate-biased diet (Table S10, p = 0.001). Note that on day 14, the area on these two diets was still symmetrical, but total area covered had not yet reached the same extent as on the optimal diet 1:4 in the no-choice experiment (Fig. 2*B*; Table S9; p = 0.16). This result showed that, by different growth patterns, the fungus regulated both amino acid and carbohydrate absorption to maximize spore production, on any of the tested complementary diet pairings, revealing its ability to solve complex nutritional challenges.

**Figure 2.**
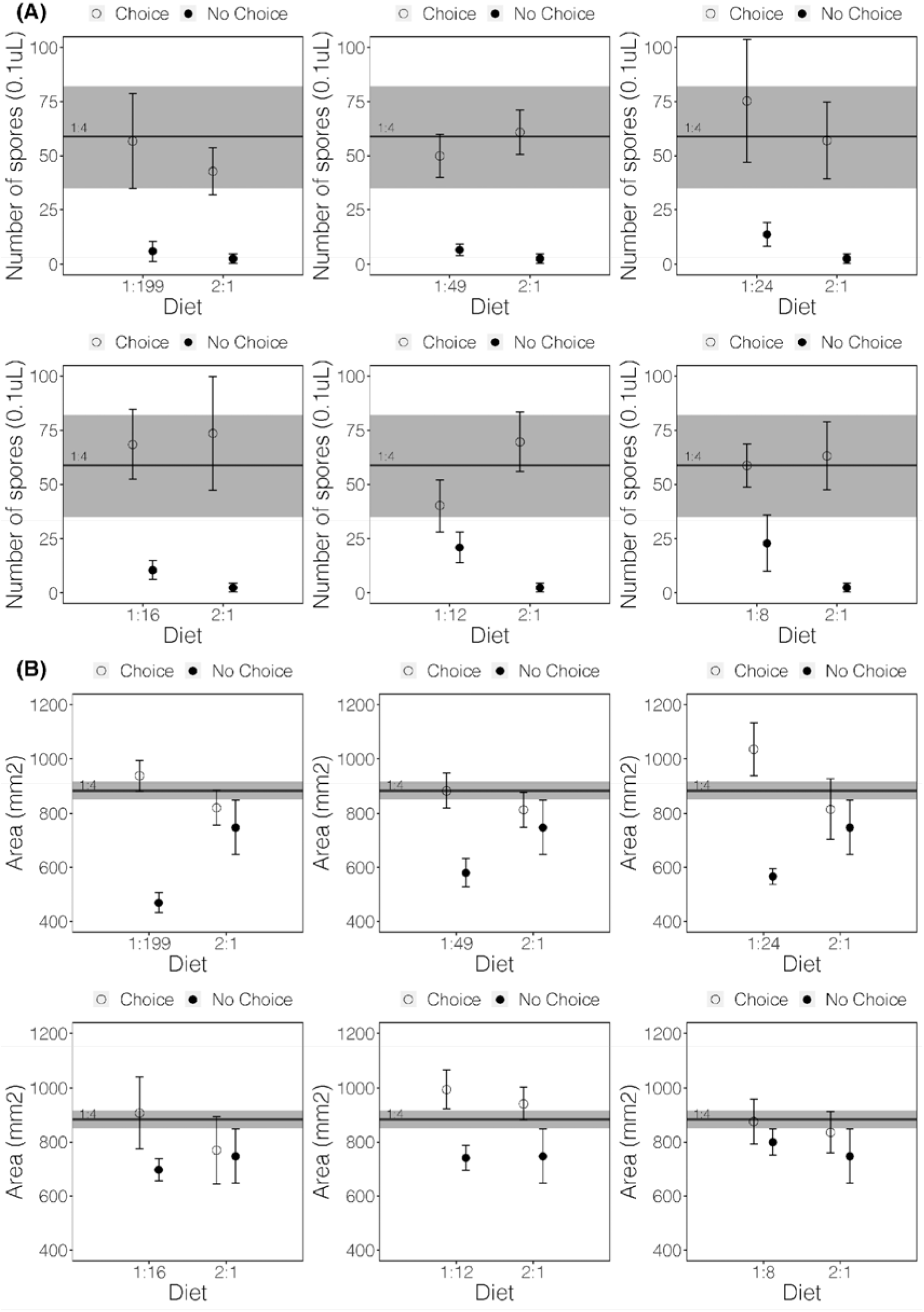
Spore production and fungal growth of *Metarhizium brunneum* under presence or absence of dietary choice (artificial diets, Table S2). **A.** Number of spores per µL (volume = 0.1µL) produced by *M. brunneum* grown for 30 days on 6 different diet pairings offering one of either C-biased diet (1:8, 1:12, 1:16, 1:24, 1:49, or 1:199) versus an AA-biased diet (2:1). For the experiment schematic design see Figure S7. For each plate, the spores were counted separately for the two diet-type halves. **B.** Fungal growth surface day 14 and **C.** day 30. The area covered by the fungus was recorded for fungi grown for 14 days and 30 days to 6 different diet pairings offering a C-biased diet (1:8, 1:12, 1:16, 1:24, 1:49, or 1:199) versus an AA-biased diet (2:1). The area was measured on each diet for each pairing. In **A, B** the black line corresponds to the results from the no-choice experiments for 1:4, the most optimal fungal diet. The grey area represents the 95% confidence interval for the 1:4 diet.

### Only healthy *Linepithema humile* ants suffer from a high-protein diet

We then identified how the survival of healthy vs *Metarhizium*-infected ants was affected by being confined to a 1:199 and 1:4 diet. Based on previous reports on ants [12, 44] we expected that the survival of uninfected *L. humile* workers would benefit from high carbohydrate levels (1:199 diet). For the infected ants, the high protein diet (1:4), being the ratio maximizing fungal performance, may allow the pathogen to replicate in the host and reduce its lifespan even further. Following the survival of a total of 200 ants for 64 days, which were kept solitarily under either the 1:199 or the 1:4 AA:C ratio diet, after being exposed to *M. brunneum* or not, showed that both diet ratio (coxph, LR χ^2^ = 6.4, p = 0.01) and infection (coxph, LR χ^2^ = 36.18, p = 0.0001) influenced the survival of the ants (with a non-significant interaction between the two (p = 0.10; Fig. 3; Table S11). As expected, infected ants died significantly faster compared to uninfected control ants on both diets. A high-carbohydrate diet (1:199) increased worker survivorship in uninfected ants (Table S12, coxph, z = −5.32, p = 0.02), whereas the survival of infected ants did not differ significantly between the two diets (Table S12, z = 0.82, p = 0.80). Importantly, this means that keeping the ants on a diet that is the optimal diet for the fungal pathogen did not increase their mortality.

**Figure 3.**
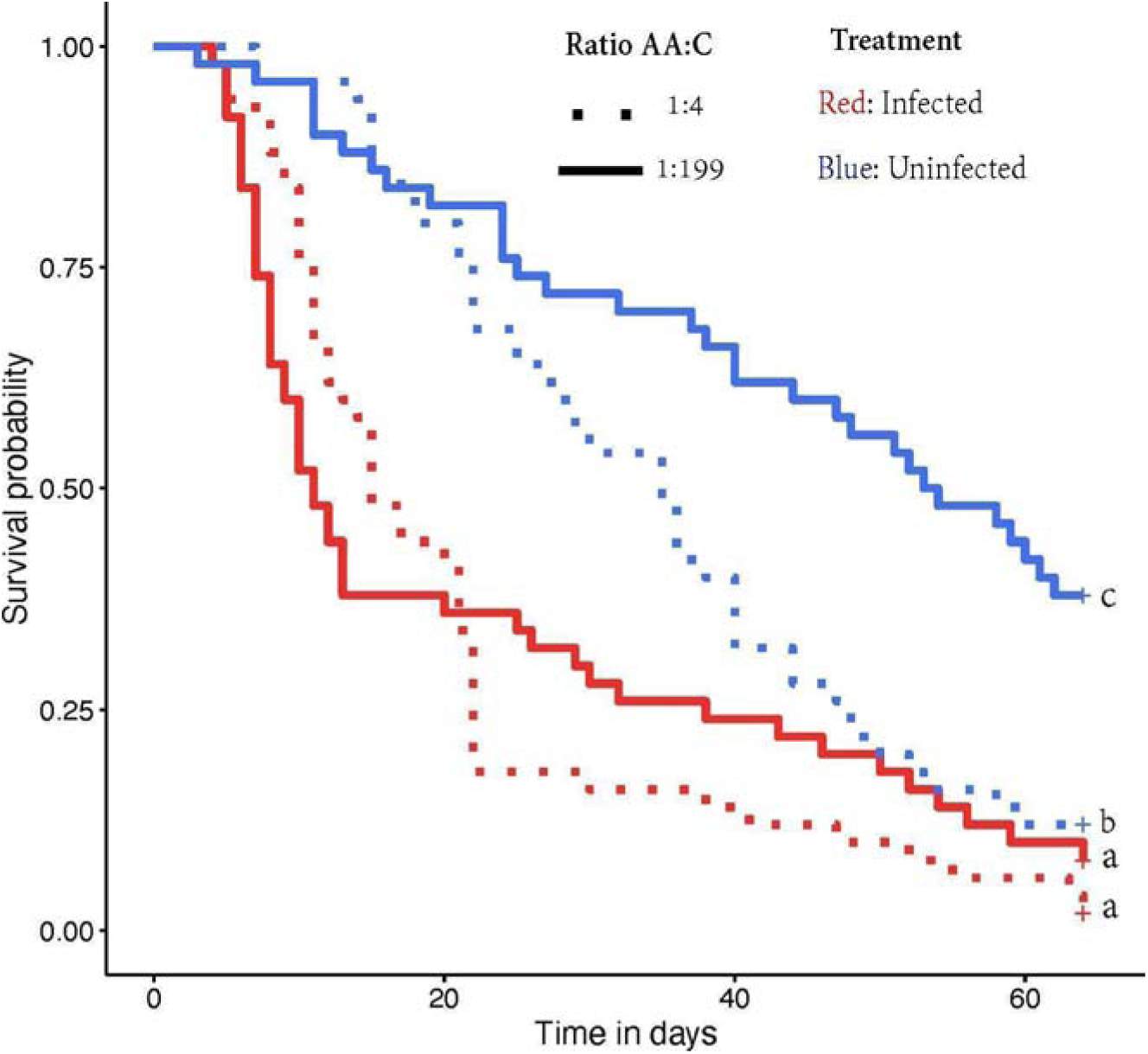
The effect of diet and infection on ant lifespan. Individual *L. humile* workers were challenged with 0.5 µl of 1 x 10^7^ spores ml^−1^ suspension of *M. brunneum*, while control groups received Triton X solution. Ants from both groups were confined to either a 1:199 or a 1:4 diet. Estimated functions for Cox regression of survival time based on the infection status and diet of *L. humile* workers.

### Infected colonies alter their preferred food choice to high-protein levels

We next tested whether infection would alter the collective food choice of the colony’s foragers, by offering both infected and uninfected colonies a choice between the carbohydrate-rich 1:199 and the more AA-rich 1:4 diet once a week over a total of 5 weeks. Foraging activity (total number of ants visiting both diets) decreased over the course of the experiment (Fig. 4*A*; Table S13, week effect p < 0.001), but was not affected by the colony’s infection status (Fig. 4*A*; Table S13, p = 0.61), despite the fact that infected colonies had higher mortality than the uninfected ones (Fig. 4*C*; Table S16, p = 0.001). Notably, however, infection altered the intake target of the colonies. Uninfected colonies selected the carbohydrate-rich 1:199 diet, which prolongs their lifespan compared to the 1:4 diet (Fig. 3). The foragers of infected colonies, however, preferentially targeted the 1:4 diet (Fig. 4*A*; Table S14, p < 0.001). Therefore, colonies regulated their food intake from a carbohydrate-biased food preference when healthy to a greatly increased intake of amino acids when infected (Fig. 4*B*; Table S15, p = 0.002).

**Figure 4.**
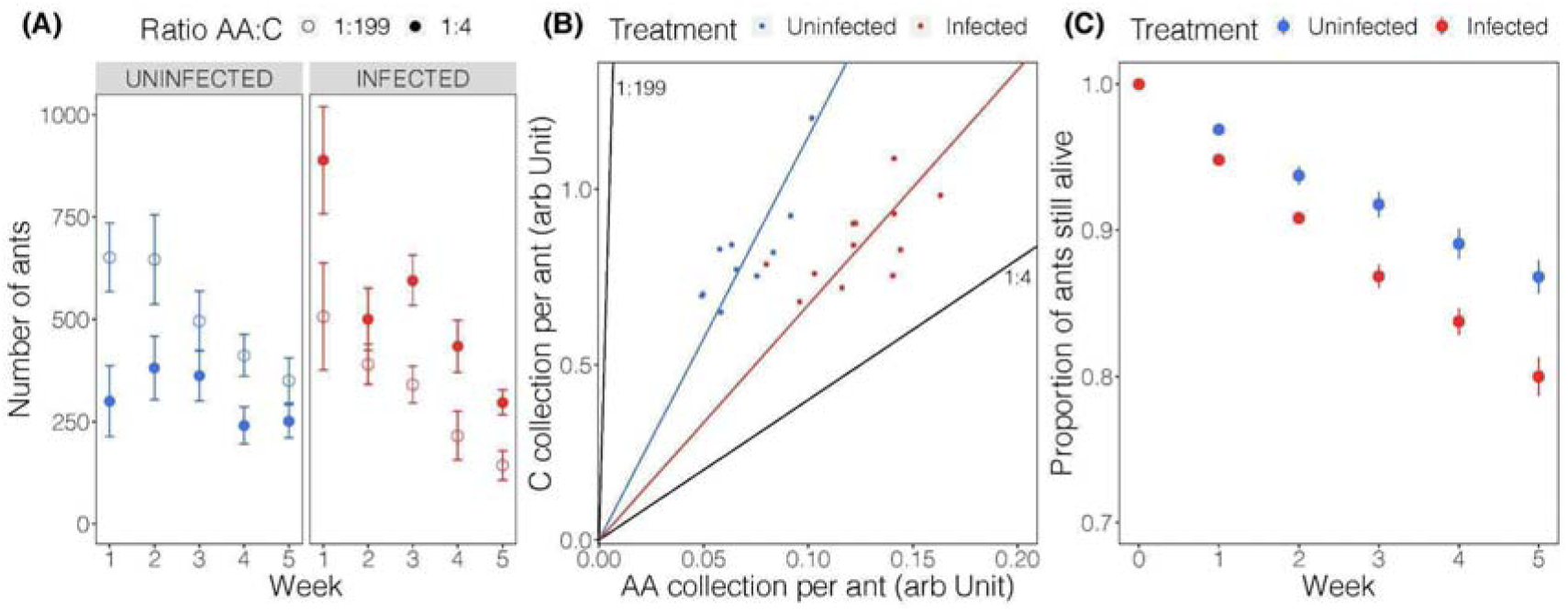
Food choice of foragers of infected and uninfected colonies. **A.** Number of foraging workers from uninfected (blue) or infected (red) colonies visiting the two diets (open circle = 1:199 and filled circle = 1:4) offered simultaneously in the food choice experiment over the course of the 5 week-period (duration of each feeding session 1 h). **B.** Amino acid (AA) and carbohydrate (C) collection during choice experiment. For each assay, the approximate nutrient collection for the colony was derived from the foraging effort (i.e., the number of ants visiting each diet) multiplied by the nutrient concentration. Points along each trajectory represent the average intake of AA and C per ant over the weeks per tested colony. Black lines show the diets 1:199 and 1:4. **C.** Proportion of infected and uninfected (control) ants alive over the 5 weeks of the experiment. Dots represent mean and error bars show 95% confidence intervals (N = 10 and 12 colonies respectively for uninfected and infected colonies). In each graph, blue color represents uninfected ants, red *Metarhizium*-infected ants.

### Immune activation induces food choice change to protein-rich diets

To disentangle, whether the choice for a higher-amino acid diet observed in infected colonies reflects a pathogen manipulation towards the fungal optimal diet (Fig. 1) or a host strategy to compensate for the increased use of amino-acids during the immune response [25,51], we subjected the ants to either a full infection with live *M. brunneum* or only induced an antifungal immune response in the absence of live pathogen by injection of fungal cell wall material (β-1,3 glucan, zymosan) – a known elicitor of the ant immune system [27,28]. Then, we compared their immune gene upregulation and food choice compared to their respective controls (Triton X-treated, resp. saline-injected ants). Immune gene expression analysis confirmed that both, *Metarhizium-*infection (p = 0.01) and β-1,3 glucan injected (p = 0.001), led to significant upregulation of antimicrobial peptides, in particular, *Hymenoptaecin* ([61], Fig. 5*A*; for other immune genes see Table S17). The ants of the four groups were then given the choice between the two diets of AA:C of 1:199 or 1:4. Interestingly, not only the *Metarhizium*-infected ants (p = 0.04; Fig. 5*B*, Table S18), but also the merely immune-activated ants (p = 0.009; Fig. 5*B*, Table S18) showed a clear upregulation of amino acid intake compared to their respective controls, and with no difference in food preferences between the *Metarhizium*-infected and β-1,3 glucan injected ants (p = 0.48; Fig. 5*B*, Table S18). This experiment unequivocally revealed that immune activation *per se*, even in the absence of a live fungal pathogen infecting the host, is sufficient to induce the change in food intake towards higher amino acid levels in the ants, in line with a compensatory mechanism for increased needs after antimicrobial peptide production during the course of the immune response.

**Figure 5.**
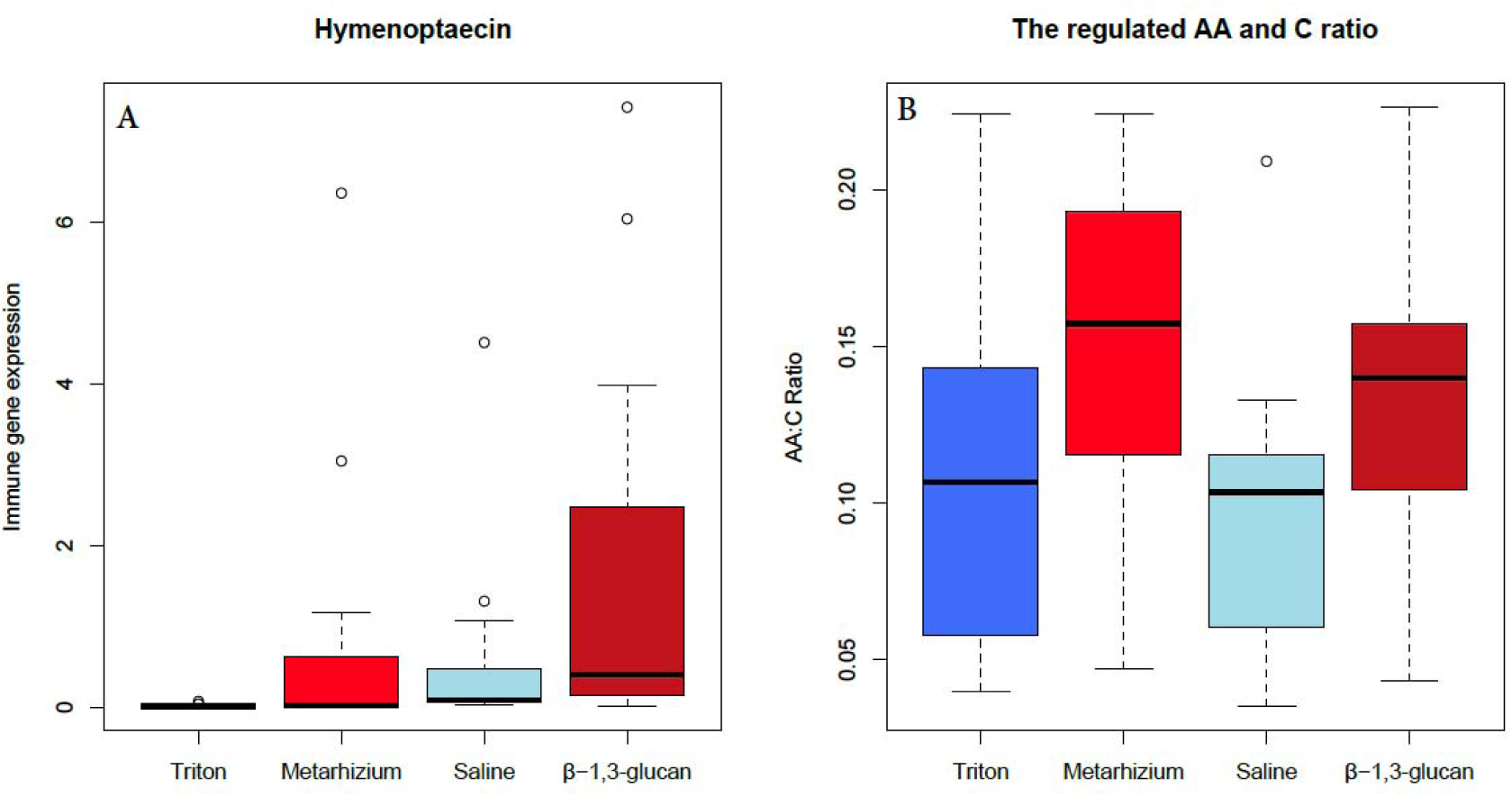
A. Immune gene expression. Candidate gene approach was applied to test if the β−1,3-glucan injection elicits an immune response in ants. The expression level of *Hymenoptaecin* (insect immune gene) was measured by droplet digital PCR (ddPCR). Expression levels of the immune gene was normalized to the housekeeping gene (*GAPDH*) before further statistical analysis. **B. The regulated amino acid (AA) and carbohydrate (C) ratio of ants (*i*) Triton-X treated, (*ii*) *Metarhizium*-infected, (*iii*) microinjected with physiological saline solution (i*v*) microinjected with** β**−1,3-glucan.** The ants had simultaneous access to both diets 1:199 and 1:4 for 1 h. Boxplots show median (black line), 25–75% quartiles, maximum/minimum range, and outliers. Dark blue represents Triton-X treated ants, light red *Metarhizium*-infected ants, light blue saline microinjected ants, while dark red β−1,3-glucan microinjected ants.

## Discussion

Using Nutritional Geometry (NG [26]), our study defines for the first time the nutritional requirements of both partners of a host-pathogen system, using social ants (*Linepithema humile*) and a fungal pathogen (*Metarhizium brunneum*). We established that the ant host and its fungal pathogen have different nutritional requirements. The optimal diet composition (intake target) of the healthy host is drastically more carbohydrate-biased than that of its pathogen. In nature, the fungus is principally feeding on hemolymph and other host tissues [29,30]. In insects, the protein-to-carbohydrate ratio of the hemolymph varies with nutritional state but also within species [31], with a reported 1:1 in silkworms to 1:6 in crickets [32,33]. A past study showed that the P:C ratio of fire ants is 1:3 [34]. Hence, the hemolymph composition of ants would appear to correspond to the optimal artificial diet composition for the pathogen, as determined in our study, making the ants a good host for this generalist entomopathogenic fungus [35–37].

In our experiments, we found the optimal diet composition of the entomopathogenic fungus *Metarhizium brunneum* in terms of growth and reproduction to be 1 to 4 essential amino acids (AA) to carbohydrate (C) ratio. Furthermore, the pathogen was able to maintain optimal growth and spore production on various complementary food pairings. Hence this fungal pathogen that comprises distributed hyphae without specialized centres is able to acquire an optimal supply of multiple nutrients essential for its reproduction on patches of different nutrient quality. Nutritional regulation has been reported only once before in a non-neural system [38].

Entomopathogenic fungi, such as *M. brunneum,* penetrate the cuticle of the host with the help of several enzymes [39–41], and then the fungus spreads into the hemolymph and kills the host within several days due to tissue penetration, toxins and nutrient depletion [39–42]. Death occurs between 4 and 10 days, depending on the virulence of the fungi and on the host species [3]. In our experiment, when confined to different diets and exposed to the fungus, *Linepithema humile* ant workers died significantly faster compared to uninfected ones on both diets (high amino acids and high carbohydrate). As found in many studies in insects and mammals [43–47], protein-rich diets often reduce lifespan, and particularly adult ants typically thrive better under higher carbohydrate diets [12,44]. This is also what we found in our experiment, where the uninfected ants survived much better under the high carbohydrate diet (1:199) than the more amino acids-rich 1:4 diet. Importantly, as observed in a previous study [12], the survival benefit of the high carbohydrate diet faded under *Metarhizium* infection, so the survival rate in infected ants did not differ significantly under the different diets (1:199, 1:4). The effects of infection were more pronounced for ants reared on 1:199, where the ants’ half-life decreased by 5-fold in response to infection, while it decreased by only 2-fold for ants reared on 1:4. Hence, we can postulate that the typical negative effects of an amino acids-richer diet must have been compensated by benefits that this diet has for infected ants. Thus, interestingly, when reared under a diet composition that corresponded to the intake target of the fungal pathogen, and not of the healthy host, the infected hosts gained a benefit, whilst one may have expected the fungus to thrive better. Our study suggests that increasing protein intake could provide a survival advantage under *Metarhizium* infection. Hence, protein intake seems to be a limiting factor that the ants should – if possible – try to increase under infection.

To see if ants would not only benefit from higher amino acids intake but would even actively choose a diet containing these nutrients, we offered infected and uninfected colonies a food choice between the diet that maximized their lifespan under healthy conditions (1:199) and the diet that improved their survivorship under infection (1:4). We found a significant difference in the nutrient ratio selected by infected and uninfected colonies; with the foragers of infected colonies choosing the 1:4 diet that under healthy conditions would reduce their survival. This change in foraging behavior could be the result of parasite manipulation, as other pathogenic fungi are capable of generating sophisticated behavioral modification in their hosts. For instance, to improve spore dispersal, *Cordyceps* and *Pandora* fungi change the ants’ behavior by promoting them to leave the nest and bite on vegetation overhanging the hosts’ foraging trail [16,48,49]. Despite intense studies [29,39,50], to our knowledge, no contribution to date has shown that any of the *Metarhizium* species is able to manipulate their hosts. Moreover, even if the 1:4 diet corresponds to the intake target of the fungus, we showed that confinement of the host on this diet did not worsen the infection, on the contrary, it provided a greater host survival.

Our results suggest that the altered foraging behavior constitutes an active response to infection of the colony, similar to solitary animals often altering their food choices following infection thereby increasing immune function [13,14,51]. In lepidopteran larvae, for example, high P:C diets lead to significantly higher levels of constitutive immune function than those on low P:C diets [13], and protein is the most important predictor of the functional (physiological) immune response [51]. This may be caused by important cellular immune defense mechanisms in insects, like the encapsulation, and melanization responses [52–54], as well as the very important effectors of the antimicrobial peptides (AMPs), which are short proteins, i.e., they require amino acids for their production [55]. Excitingly, in our collective choice experiment, we could observe this change in strategy in the foragers, even if only 0.2% of colony members died from infection. This reveals that foraging individuals were able to very sensitively react to the altered needs of a minority of colony members, thereby provisioning the colony with a diet composition that helped to keep infections largely at bay. Our findings can therefore be compared to self-medication in solitary species, where uptake of otherwise harmful diets provides a benefit under disease [solitary insects: 56,57 ants: 58,59].

To demonstrate that infected ants chose a higher amino acid diet to recover the resources used up during the immune response and not as a result of pathogen manipulation, we injected ant workers with β-1,3 glucan to induce an immune response in the absence of an actual pathogen [60]. We could confirm, that indeed, injection of β-1,3 glucan but not saline (control) caused an increase in the expression of the immune gene, namely, *Hymenoptaecin,* which is a glycine-rich antimicrobial peptide [55,61], and requires amino acids for its production [55]. Our results showed that the chosen AA:C ratio was more protein-biased for β-1,3 glucan-injected and infected ants than their respective controls suggesting a compensatory strategy to compensate for the amino acids costs of immune activation. In summary, we found that the change in foraging behavior observed in infected colonies could be a protective behavior used by the colony to combat an infection, a potential process of “collective self-medication”. This adaptive foraging strategy suggests a novel social immunity mechanism to promote colony-level health [62,63]. Our work paves the way for more studies investigating nutritional immunology at the social level.

## STAR Methods

### KEY RESOURCES TABLE

| REAGENT or RESOURCE | SOURCE | IDENTIFIER |
| --- | --- | --- |
| **Chemicals, peptides, and recombinant proteins** |  |  |
| Saccharose | Sigma-Aldrich | CAS#57-50-1 |
| Ethanol | Fisher scientific | CAS#64-17-5 |
| Cholesterol | Sigma-Aldrich | CAS#57-88-5 |
| Choline Chloride | Sigma-Aldrich | CAS#67-48-1 |
| Acid Ascorbic | Sigma-Aldrich | CAS#50-81-7 |
| Vanderzant Vitamin Mix | Fisher scientific | CAT#903244 |
| Wesson Salt Mix | Fisher scientific | CAT#902851 |
| Inositol | Sigma-Aldrich | CAS#87-89-8 |
| Methionine | Sigma-Aldrich | CAS#63-68-3 |
| Histidine | Sigma Aldrich | CAS#71-00-1 |
| Valine | Sigma-Aldrich | CAS#72-18-4 |
| Threonine | Sigma-Aldrich | CAS#72-19-5 |
| Arginine | Sigma-Aldrich | CAS#74-79-3 |
| Tryptophan | Sigma-Aldrich | CAS#73-22-3 |
| Phenylalanine | Sigma-Aldrich | CAS#63-91-2 |
| Leucine | Sigma-Aldrich | CAS#61-90-5 |
| Isoleucine | Sigma-Aldrich | CAS#73-32-5 |
| Lysine | Sigma-Aldrich | CAS#56-87-1 |
| Fluon | Whitford | U-22101540 |
| Malt Extract Agar | Sigma-Aldrich | 70145 |
| Triton™ X-100 | Sigma-Aldrich | 1056142500 |
| Sodium hypochlorite | Sigma-Aldrich | CAS#9002-93-1 |
| Physiological ant saline solution |  |  |
| Zymosan-A | Sigma-Aldrich | CAS#58856-83-2 |
| Serpin 27a | Sigma-Aldrich |  |
| Defensin | Sigma-Aldrich |  |
| Relish | Sigma-Aldrich |  |
| Hymenoptaecin | Sigma-Aldrich |  |
| **Deposited data** |  |  |
| All Data | Supplementary Material |  |
| Code | Supplementary Material | Data S1 |
| **Experimental models: Organisms/strains** |  |  |
| *Metarhizium brunneum* (strain Ma 275, KVL 03-143) |  |  |
| Argentine ants *Linepithema humile* | Université Paul Sabatier<br>- Toulouse | N/A |
| **Software and algorithms** |  |  |
| R Core Team 2021 | A language and environment for statistical computing. R Foundation for Statistical Computing, Vienna, Austria. | URL<br>https://www.R-project.org/. |
| Antethoc | Author: Maud Combe | https://cran.r-project.org/web/packages/lme4/index.html |
| ImageJ | Schneider et al., 2012 | https://imagej.nih.gov/ij/ |

### Lead contact

Further information and requests for resources should be directed to and will be fulfilled by the **Lead Contact** Enikő Csata.

### Materials availability

The study did not generate new unique materials.

### Experimental models and subject details

#### Host ant

We used the Argentine ant, *Linepithema humile*, a widespread invasive species, that forms polygynous supercolonies with millions of individuals [64,65]. Argentine ants are opportunistic and feed on a large variety of nutritional resources, such as protein-rich prey items, as well as carbohydrate-rich nectar and honeydew [66]. Generally, ants within the colony have their own nutritional needs, adults such as foragers and nurses mostly need carbohydrates as a source of energy, whereas larvae and reproductive adults rely also heavily on proteins for growth and egg production respectively [67]. Ant colonies were collected from a large supercolony that lacks nest boundaries in Toulouse (France) in 2018, following the guidelines of the Nagoya protocol for Access and Benefit Sharing. Ants were subdivided into experimental colonies (see details below). When not used in experiments, we supplied each colony with water and a mixed diet of vitamin-enriched food [68]. All experiments followed European and national law and institutional guidelines.

#### Fungal pathogen and conidiospore suspension

We exposed experimental colonies of Argentine ants to the entomopathogenic fungus *Metarhizium brunneum* (strain Ma 275, KVL 03-143), previously known as *Metarhizium anisopliae*, but now separated as a sister species [35]. The genus *Metarhizium* is used worldwide for invertebrate pest control [69,70]. *M. brunneum* is a generalist entomopathogenic fungus that infects several host species from different taxa [50] and is a natural pathogen of ants [35].

### Method details

#### Experiments and measures Intake Target of the fungus *Metarhizium brunneum*

18 synthetic diets were prepared and filter-sterilized (filter: 0.22 μm polyethersulfone membrane). The diets varied in both the amino acid to carbohydrate ratio and total concentration of amino acid + carbohydrate. The amino acid content of the diets consisted of a mixture of the following essential amino acids: Arginine, Histidine, Isoleucine, Leucine, Lysine, Methionine, Phenylalanine, Threonine, Tryptophan, and Valine, whereas glucose was used as a digestible carbohydrate source. The fat content of the diet consisted of a mix of cholesterol and lecithin. Each diet contained the same quantity of fat and other nutrients (lecithin 0.034%, cholesterol 0.034%, Vanderzant Vitamin mix 0.1%, Wesson salt mix 0.1%, ascorbic acid 0.2%, inositol 0.05%, choline chloride 0.034%). All the nutrients were mixed in a 0.66% agar gel and this mixture was poured into Petri dishes (10 mL in a 55 mm x 15 mm Petri dish). The Petri dishes were then sealed with parafilm and stored at 4**°**C.

##### Experiment 1: No-choice diet experiment

In our first experiment, we investigated how the entomopathogenic fungus *Metarhizium brunneum* reaches its Intake Target (IT). The fungus *M. brunneum* was confined to 18 different diets, varying in their essential amino acids (AA) and carbohydrate (C) ratios (2:1, 1:1, 1:1.5, 1:2, 1:3, 1:4, 1:6, 1:8, 1:12, 1:16, 1:24, 1:49, 1:99, 1:199) with other essential ingredients (*see* Table S1). The diets were poured into Petri dishes (10 mL in a 55 x 15 mm Petri dish). For each diet, 10 µl of a 2 x10^7^ spores ml^−1^ suspension was applied in the centre of a Petri dish directly on the diet, and then the Petri dish was sealed with parafilm and incubated at 25**°**C.

##### Experiment 2: Dilution experiment

In the second experiment, we investigated whether the entomopathogenic fungus expands a greater area on more diluted diets, thus increasing the surface in contact with the diet and compensating for nutrient dilution (*see* Table S2). The diets were poured (10 mL) into Petri dishes (55 x 15 mm). For each diet, 10 µl of a 2 x 10^7^ spores ml^−1^ suspension were applied in the centre of a Petri dish directly on the diet, then sealed with parafilm and incubated at 25°C. The following diets were diluted: 1:4 (control 1:4, 1:4/2, 1:4/4, 1:4/8, 1:4/x2).

For the first and the second experiment, the fungal growth was recorded using a camera (Lumix DMC-FZ1000) after 14 and 30 days. The fungal expansion (growth rate: area mm^2^) was measured using the program ImageJ (NIH Image, v1.49g). The fungus was harvested after 30 days from each diet by taking three subsamples of 1.5 cm diameter at equal distances from the centre of the dish, and the number of spores was counted from each subsample. The fungal growth never reached the edge of the Petri dishes in the different experiments (*no-choice* and *choice*). Spores were counted with a hemocytometer (depth 0.1 mm, Sigma-Aldrich, catalog number: Z359629-1EA) and an automatic cell counter (Nexcelom Bioscience, Cellometer Auto M10). The relationship between the two counting methods (hemocytometer and cellometer) were significant (R-squared = 0.67, p **<** 0.001, N = 55).

##### Experiment 3: The effect of past diet on spores production

In our third experiment, we tested the capacity of the spores produced on different diets for growth and spore production when they were all grown on a standard rearing medium (Malt Extract Agar). We harvested spores from the two most extreme diets (1:199 and 2:1) and from the balanced one 1:4. Then, 10 µl of 30 spores ml^−1^ suspension were applied in the center of a Petri dish (diameter 55 mm) directly on a standard rearing medium (10 mL Malt Extract Agar medium: agar 1.5%, malt extract 3%, mycological peptone 0.5%). We replicated the experiment at least 5 times for each. Then, the Petri dishes were sealed with parafilm and incubated at 25**°**C. The fungus was harvested after 30 days. Spores were harvested from three subsamples of 1.5 cm diameter from each Petri dish. Spores were counted with a hemocytometer (depth 0.1 mm, Sigma-Aldrich, catalog number: Z359629-1EA).

##### Experiment 4: Choice diet experiment

In our fourth experiment, we allowed the fungus *M. brunneum* to select between two diets differing in their content of essential amino acids (AA) and carbohydrate (C) ratio. Following the results from the first experiment, we tested six diet pairings (1:199 vs. 2:1, 1:49 vs. 2.1, 1:24 vs. 2:1, 1:16 vs. 2:1, 1:12 vs. 2:1, 1:8 vs. 2:1). The Petri dishes (diameter 55 mm) were divided into two parts, one of the diets was poured on one side of the Petri dish (5 mL) and the second diet was poured on the other side of the Petri dish (5 mL). The second diet was poured after the first solidified. To each Petri dish, 20 µl of a 2 x 10^7^ spores ml^−1^ suspension was applied in a line, in the center of a Petri dish directly in contact with both diets (55 x 15 mm, Fig. S7). The Petri dishes were sealed with parafilm and incubated at 25**°**C. The fungal growth was recorded with a camera (Lumix DMC-FZ1000) after 14 and 30 days. The fungal expansion (growth rate: area mm^2^) was measured using the program ImageJ (NIH Image, v1.49g). The fungus was harvested after 30 days from each diet by taking two subsamples from each diet, and the number of spores was counted for each subsample. Spores were counted with a hemocytometer (depth 0.1 mm, Sigma-Aldrich, catalog number: Z359629-1EA).

#### Intake Target of the host ant: *Linepithema humile*

##### Experiment 5: Survival of the host on different diets

For survival assays, we studied how infected and uninfected hosts performed when confined to diets offering varying amounts of amino acids and carbohydrates. Ants were confined to two different diets, varying in their essential amino acids (AA) and carbohydrate (C) ratios (1:199, 1:4). Then, ants were infected with *Metarhizium* spores, suspended in 0.05% Triton-X. We applied 0.5 µl of 1 x 10^7^ spores ml^−1^ suspension to the ants. Control ants received 0.5 µl of 0.05% Triton-X solution. In total 200 ants (100 infected and 100 control) were housed solitary, in Fluon-coated Petri dishes (55 x 15 mm) with moistened cotton (remoistened every day). The mortality of ants was checked daily, for over 64 days. We supplied ants with a diet three times per week. Dead ants were removed from the nest, surface sterilized, and monitored for 14 days for signs of *Metarhizium* infection. At the end of the experiment (64 days), the live ants were killed, surface sterilized, and monitored for signs of *Metarhizium* infection. In total 200 infected and uninfected corpses were surface-sterilized. We observed that 65.26% of the 100 corpses from infected groups and none of the 100 corpses from control groups produced *Metarhizium* conidia. In *Metarhizium*-exposed ants, the proportion of cadavers sporulating did not differ significantly with diet (GLM, χ^2^ = 0.73, p = 0.39, high carbohydrate diet (1:199) = 63.04%, high essential amino acids diet (1:4) = 67.3%).

##### Experiment 6: Collective choice of infected and uninfected ants

This experiment allowed us to determine whether when given a choice, ants may actively select a diet in response to the colony’s health state. Colonies of uninfected and infected *L. humile* were offered two diets varying in the ratio of essential amino acids (AA) and carbohydrate (C). Ants were offered a solution of 1:199 AA:C or a solution of 1:4 AA:C for one hour. We investigated whether ants preferred 1:199 or a solution of 1:4, or whether they showed no preference depending on their infection state. We used 22 colonies in total, 12 were infected while 10 were uninfected. Each colony contained 2000 individuals, one queen and brood.

For each experimental colony, ants were installed in 3 test tube nests (15 cm in length, 1.3 cm in diameter). These tubes were placed in a rearing box (20 x 10 x 10 cm) with walls coated with Fluon to prevent ants from escaping. Between the choice assays, ant colonies were food-deprived and kept at room temperature (25 ± 1°C) under a 14:10 L:D photoperiod. To infect experimental colonies with *M. brunneum*, spores were diluted in 0.05% Triton solution (1 x 10^7^ spores per mL). Then, 450 µl of this suspension was vortexed vigorously and deposited on a filter paper (7 cm diameter) which was placed in the rearing box (modified after [71]). For the control groups, 450 µl of 0.05% Triton X solution was deposited on the filter paper, following the same procedure. We renewed these operations every three days for each treatment for 5 weeks (modified after [72]). Hence, the workers were in contact with the fungal spores during the entire time course of the experiment.

Ants were starved for the whole duration of the experiment (5 weeks) except when we were running the diet choice assay once a week. The assay started 3 days after the first exposure to the fungal spores. During an assay, two diets (1:199 and 1:4) were placed on two platforms (5 cm x 5 cm) tied to a Y-shaped bridge with two branches of equal length (L = 5 cm, 60 angle between the two branches) connected to the colony. For each assay, we added a colorant randomly to each solution and each examiner was blinded to the identity of the solution tested. The ants had access to both solutions for one hour [73]. The ant traffic on the two branches was recorded by a video camera (Canon LEGRIA HF G30) for one hour. To assess foraging effort, for all choice assays, we counted the number of ants travelling on each branch at a particular point (one centimeter from the choice point) every minute for one hour. Counting began as soon as the first ant climbed onto the bridge and lasted for 60 min [73]. During the entire experiment, we also counted the number of dead ants once a week. Dead ants were surface-sterilized and monitored for 14 days for signs of *Metarhizium*-infection. During the experiment, in total 625 infected and 485 uninfected ants died and their corpses were surface-sterilized and observed for fungal outgrowth. We found that 11.68% of the 625 corpses from infected groups and none of the 485 corpses from control groups produced *Metarhizium* spores. These last results prove that our infection method that mimics natural conditions worked successfully.

##### Experiment 7: Host compensation or pathogen manipulation

To test whether the changes observed in the collective foraging behavior of infected ants are caused by host compensation or pathogen manipulation, we stimulated the ant’s immune system using β-1,3 glucan. If immune-challenged ants forage mostly on the 1:4 AA:C diet similarly to infected ants this would suggest that ants are using AA to fight the infection. On the contrary, if they forage mostly on the 1:199 AA:C diet as uninfected ants, this would suggest that infected ants are responding to parasite manipulation. ***<u>Immune gene</u> <u>expression analysis.</u>*** A candidate gene approach was applied to test if the β-1,3 glucan injection elicited an immune response in ants, using saline-injected ants as controls. We also quantified the immune response of *Metarhizium*-infected ants using Triton-X-treated ants as controls. We measured the expression levels of four immune genes commonly found in insects, including ants (*Defensin 1*, *Hymenoptaecin*, *Relish*, *Serpin 27a*) by droplet digital PCR (ddPCR). *Defensin 1* and *Hymenoptaecin* are genes coding for antimicrobial peptides [61, 76]; *Relish* is required for humoral immune response [77]; and *Serpin 27a* (SPN-27a) a gene involved in the melanization cascade [78] and *GAPDH* as a housekeeping gene. Sample preparation and gene expression analyses were performed as detailed in the electronic supplementary material. Expression levels of the immune genes were normalized to the housekeeping gene (*GAPDH*) before further statistical analysis (for primer sequences see Table S19). Soluble β-1,3 glucan was required by suspending 5 mg of Zymosan-A (*Saccharomyces cerevisiae* wall fragments [Sigma-Aldrich]) in 1 ml of sterile physiological ant saline (as described in [74]). The zymosan suspension was vortexed for 1 hr at 3200 rpm before being centrifuged at 10000 rcf for 5 min. The supernatant, containing the soluble

β-1,3 glucan [75] was then removed and stored at 4°C. Ant individuals were placed gently into a sponge harness. Using fine glass capillaries (with a spike to aid injection; inner diameter = 25 µm [BioMedical Instruments, Germany]), a microinjector (parameters: pi = 120 hPa, ti = 0.3 s, pc = 20 hPa [FemtoJet, Eppendorf, Germany]) and a micromanipulator (Luigs and Neumann, Germany) we injected 46 nl of the β-1,3 glucan solution (N = 21) through the intersegmental membrane of ants’s first tergite, into their haemocoel. Capillaries were cleaned between injections using 96% ethanol. As a control, ants were microinjected with sterile physiological saline (N = 15, [74]). Other groups of ants were infected with *Metarhizium* spores (10^7^ spores ml^−1^ suspension, N = 16), and control ants received 0.5 µl of 0.05% Triton-X solution (N = 16, for further details *see* Supplementary methods). ***<u>Individual choice assay</u>***. Before each choice assay, individuals were food-deprived and isolated for 24 hours. After each treatment (*i*) microinjected β-1,3 glucan, *ii*) microinjected with physiological saline solution, (*iii*) *Metarhizium*-infected (i*v*) Triton X-treated ants were housed qindividually in Petri dishes (55 x 15 mm) and were food-deprived. After 24 hours, ant individuals were offered two diets varying in the ratio of essential amino acids (1:4 AA) to carbohydrate (1:199 C). Individuals were filmed with Ueye cameras for 1 hr (whereby 4 cameras were used in parallel, each filming 2 replicates simultaneously). Just after the experiment, individuals were frozen at – 80°C for the analysis of their immune gene expression.

## Data and code availability

- All data generated in this paper will be made publicly available in a data repository such as Dryad upon acceptance.
- Any additional information required to reanalyze the data reported in this paper is available from the lead contact upon request.

## Acknowledgements

We are thankful to Jorgen Eilenberg and Nicolai V. Meyling for the fungal strain, to Simon Tragust, Abel Bernadou, and Brian Lazarro for insightful discussions, to Iago Sanmartín-Villar, Léa Briard, Céline Maitrel, and Nolwenn Rissen for their help during experiments. Furthermore, we thank Anna Grasse for the help with the immune gene expression analyses. E.C. was supported by the Fyssen Foundation Grant and the Alexander von Humboldt Foundation. A.D. was supported by the CNRS.

## Author Contributions

A.D. and E.C. conceived the study. Experiments were designed by E.C., A.D and S.C., and performed by E.C., E.L. and H.L., using resources of the laboratories of A.D., S.C. and J.H. Data were curated by E.C., and statistically analyzed and designed as figures by E.C. and A.D. with input of A.P.E., S.J.S. The manuscript was written by E.C., A.D., A.P.E., S.C., J.H. and SJ.S. All authors approved the manuscript.

