## Supplementary material for "Fungal infection alters collective nutritional intake of ant colonies": Csata_et_al_2023_Supplementary.docx

**Supplementary Methods**

Conidiospore suspension and viability

The fungal conidiospores (hereafter abbreviated as spores) was harvested from Malt Extract Agar (MEA) plates using 10 mL of sterile 0.05% Triton X-100, after a minimum of three weeks of growth and were washed by centrifuging them for 5 minutes at 3000 g. The supernatant was discarded and the pellet was resuspended in sterile 0.05 % Triton X. This step was repeated three times. A hemocytometer (depth 0.1 mm, Sigma-Aldrich, catalog number: Z359629-1EA) was used to estimate the concentration of the conidial suspension, which was stored at 4**°**C.

The viability of conidia was checked before each experiment by plating them on MEA plates, which were parafilmed, placed upside-down and incubated at 25**°**C overnight. The plates were checked the next day for hyphal growth. The germination of conidia was > 97% for all experiments. To confirm successful *Metarhizium* infections of the ants, we checked them after their death for fungal outgrowth and sporulation. Ant corpses were first surface-sterilized by washing in 70% ethanol for ten seconds and then in distilled water for a few seconds. Following this first rinse, ants were placed in diluted sodium hypochlorite (0.5 % NaCl) for one min, rinsed twice briefly in distilled water and subsequently dried on filter paper for a few minutes (1). Lastly, the corpses were transferred to closed Petri dishes on a moistened filter paper with a cotton wool ball soaked in water and placed in the middle of the Petri dish. The Petri dishes were kept in an incubator for 14 days at 23**°**C, and the sporulation of the fungus was checked every day for 2 weeks. The same protocol was used for the control groups (uninfected ants). No fungal outgrowth was observed in the case of uninfected ants.

Immune gene expression by ddPCR

The samples (n = 68) were homogenized using a TissueLyser II (Qiagen, 85300) with a mixture of one 2.8 mm ceramic (VWR, MOBO13114-325), five 1 mm zirconia beads (BioSpec Products; Lactan, N0381) and approx. 100 mg of 425–600 μm glass beads (Sigma-Aldrich, G8772) in 50 μl water. Total DNA was extracted using the DNeasy blood and tissue kit (Qiagen, 69506) according to the manufacturer's recommendations, with a final elution volume of 50 µl buffer AE. The genomic DNA was digested using EcoRI-HF and HindIII-HF enzymes (both New England Biolabs, R3101S and R3104S) within the 20 μl 1x ddPCR reaction, comprising: 10 μl 2× ddPCR Supermix for probes (Bio-Rad, 1863010), 18 pmol of both PR1 primers (forward primer and reverse primer as given above for the qPCR; Sigma-Aldrich), 5 pmol of the *M. robertsii* PR1 probe (as above, yet labelled with 6FAM), 10 U each of EcoRI-HF and HindIII-HF, 4.8 μl nuclease-free water (Sigma-Aldrich, W4502-IL) and 2 μl DNA template. Droplet generation was done using the QX200 droplet generator (Bio-Rad, 17005227) according to the manufacturer’s instructions. Droplets were transferred into a 96-well plate (Eppendorf, 0030128575) for PCR amplification in a T100 thermal cycler (Bio-Rad, 1861096). Cycling conditions were as follows: enzyme activation for 10 min at 95 °C, followed by 40 cycles of 30 s at 94 °C and 1 min at 56 °C, then enzyme deactivation for 10 min at 98 °C. For the entire protocol, the ramp rate was set to 2 °C s−1. Following PCR amplification, the PCR plate was put into a QX200 droplet reader (Bio-Rad, 1864003) for the readout of positive and negative droplets. Data analysis was done using the QuantaSoft Analysis Pro software (Bio-Rad). The thresholds were set manually to 3,500. The absolute number of spores per sample was computed from the obtained copy number per well by adjusting to the amount of template used in the ddPCR reaction (2 µl), the elution volume used during DNA extraction.

**Statistical analyses**

All analyses and statistics were conducted in R software (version 4.1.0). The nutritional response landscape for spore number per µL and area covered were evaluated with a surface regression model. We first averaged the number of spores for each Petri dish (triplicates for each Petri dish). We scaled the predictors in our model by subtracting their mean value and dividing them by the standard deviation (so their values are centered at 0 and in an interval close to [-1,1]). This procedure reduces the covariation between linear variables and their interaction terms.

The response variables that did not fit linear model requirements were transformed using the “*bestNormalize*” function, “*bestNormalize*” package (2). In our models, when interactions were non-significant, we ran the models again without this interaction and selected the best model using AIC-based model selection. In the case of Figure 1, response surfaces were visualized using nonparametric thin-plate splines, which were fitted using the “*fields*” package in the statistical software R (3).

The experimental lifespan of an individual was calculated as the number of days from the beginning of observations until its death or until the termination of the experiment (64 days). The Cox regression model (proportional hazard approach, N = 200 individuals) was used to analyze the effects of infection and diets on the survival of ants. The infection state of the ants (infected vs. uninfected) and the different diets were included in the test explanatory variables. Cox regression analysis was carried out with the use of the “*coxph*” function in the package “*survival*” (4). Tukey’s HSD tests were used to calculate post-hoc comparisons on each factor in the model using the function “*glht*” from the R package “*multcomp*” (5). The relationship between the two different spore-counting methods was analysed with a linear regression model. The effect of the different treatments (Triton X-treated, *Metarhizium-*infected, Saline-injected, and β-1,3 glucan injected) on the individual food choice was analysed using a linear mixed effect model approach (LMM). The day of the tests was included as a random factor to account for dependencies. LMMs were performed using “*lmer*” function both in “*lme4*” package (6), and “*car*” package with “*ANOVA*” function, to identify the exact significance level of input variables (7). To test whether the different treatments influenced the immune response of the ants (immune genes: Relish, Spn, Hym and Defensin), we performed a linear model approach (LM). In the models, when interactions were non-significant, we ran the models again without this interaction and selected the best model using AIC-based model selection. Expression levels of the immune genes were normalized to the housekeeping gene (*GAPDH*) before further statistical analysis. The graphs were carried out using the “*ggplot2*” R package (8).

**Supplementary Tables**

**Table S1.** **Diet composition.** Diets offered during the nutritional treatment for the ants (survival and collective choice) and also for the *Metarhizium brunneum* entomopathogenic fungus (no-choice diet experiments, choice diet experiment). Unless otherwise stated, quantities are given in grams. The following amino acids (AA) were used: M = Methionine, H = Histidine, V = Valine, R = Arginine, T = Threonine, W = Tryptophan, I = Isoleucine, L = Leucine, F = Phenylalanine, K = Lysine.

| **Diets** | | **2:1** | **1:1** | **1:1.5** | **1:2** | **1:3** | **1:4** | **1:6** | **1:8** | **1:12** | **1:16** | **1:24** | **1:49** | **1:99** | **1:199** |
| --- | --- | --- | --- | --- | --- | --- | --- | --- | --- | --- | --- | --- | --- | --- | --- |
| Sucrose | | 11.11 | 16.67 | 20 | 22.22 | 25 | 26.67 | 28.53 | 29.60 | 30.80 | 31.33 | 32 | 32.67 | 33 | 33.17 |
| AA | M | 2.22 | 1.67 | 1.33 | 1.11 | 0.83 | 0.67 | 0.48 | 0.37 | 0.25 | 0.20 | 0.13 | 0.07 | 0.03 | 0.02 |
|  | H | 2.22 | 1.67 | 1.33 | 1.11 | 0.83 | 0.67 | 0.48 | 0.37 | 0.25 | 0.20 | 0.13 | 0.07 | 0.03 | 0.02 |
|  | V | 2.22 | 1.67 | 1.33 | 1.11 | 0.83 | 0.67 | 0.48 | 0.37 | 0.25 | 0.20 | 0.13 | 0.07 | 0.03 | 0.02 |
|  | R | 2.22 | 1.67 | 1.33 | 1.11 | 0.83 | 0.67 | 0.48 | 0.37 | 0.25 | 0.20 | 0.13 | 0.07 | 0.03 | 0.02 |
|  | T | 2.22 | 1.67 | 1.33 | 1.11 | 0.83 | 0.67 | 0.48 | 0.37 | 0.25 | 0.20 | 0.13 | 0.07 | 0.03 | 0.02 |
|  | W | 2.22 | 1.67 | 1.33 | 1.11 | 0.83 | 0.67 | 0.48 | 0.37 | 0.25 | 0.20 | 0.13 | 0.07 | 0.03 | 0.02 |
|  | I | 2.22 | 1.67 | 1.33 | 1.11 | 0.83 | 0.67 | 0.48 | 0.37 | 0.25 | 0.20 | 0.13 | 0.07 | 0.03 | 0.02 |
|  | L | 2.22 | 1.67 | 1.33 | 1.11 | 0.83 | 0.67 | 0.48 | 0.37 | 0.25 | 0.20 | 0.13 | 0.07 | 0.03 | 0.02 |
|  | F | 2.22 | 1.67 | 1.33 | 1.11 | 0.83 | 0.67 | 0.48 | 0.37 | 0.25 | 0.20 | 0.13 | 0.07 | 0.03 | 0.02 |
|  | K | 2.22 | 1.67 | 1.33 | 1.11 | 0.83 | 0.67 | 0.48 | 0.37 | 0.25 | 0.20 | 0.13 | 0.07 | 0.03 | 0.02 |
| Cholesterol | | 0.17 | 0.17 | 0.17 | 0.17 | 0.17 | 0.17 | 0.17 | 0.17 | 0.17 | 0.17 | 0.17 | 0.17 | 0.17 | 0.17 |
| Lecithin | | 0.17 | 0.17 | 0.17 | 0.17 | 0.17 | 0.17 | 0.17 | 0.17 | 0.17 | 0.17 | 0.17 | 0.17 | 0.17 | 0.17 |
| Ethanol (mL) | | 5 | 5 | 5 | 5 | 5 | 5 | 5 | 5 | 5 | 5 | 5 | 5 | 5 | 5 |
| Vanderzant Vitamin Mix | | 0.5 | 0.5 | 0.5 | 0.5 | 0.5 | 0.5 | 0.5 | 0.5 | 0.5 | 0.5 | 0.5 | 0.5 | 0.5 | 0.5 |
| Wesson Salt Mix | | 0.5 | 0.5 | 0.5 | 0.5 | 0.5 | 0.5 | 0.5 | 0.5 | 0.5 | 0.5 | 0.5 | 0.5 | 0.5 | 0.5 |
| Ascorbic acid | | 1 | 1 | 1 | 1 | 1 | 1 | 1 | 1 | 1 | 1 | 1 | 1 | 1 | 1 |
| Inositol | | 0.25 | 0.25 | 0.25 | 0.25 | 0.25 | 0.25 | 0.25 | 0.25 | 0.25 | 0.25 | 0.25 | 0.25 | 0.25 | 0.25 |
| Choline chloride | | 0.17 | 0.17 | 0.17 | 0.17 | 0.17 | 0.17 | 0.17 | 0.17 | 0.17 | 0.17 | 0.17 | 0.17 | 0.17 | 0.17 |
| Agar | | 3.33 | 3.33 | 3.33 | 3.33 | 3.33 | 3.33 | 3.33 | 3.33 | 3.33 | 3.33 | 3.33 | 3.33 | 3.33 | 3.33 |
| Distilled water (mL) | | 500 | 500 | 500 | 500 | 500 | 500 | 500 | 500 | 500 | 500 | 500 | 500 | 500 | 500 |

**Table S2.** **Diet composition.** Diets offered during the nutritional treatment for the *Metarhizium brunneum* entomopathogenic fungus (dilution experiment). Unless otherwise stated, quantities are given in grams. The following amino acids (AA) were used: M = Methionine, H = Histidine, V = Valine, R = Arginine, T = Threonine, W = Tryptophan, I = Isoleucine, L = Leucine, F = Phenylalanine, K = Lysine.

| **Diets** | | **1:4** | **1:4/2** | **1:4/4** | **1:4/8** | **1:4/x2** |
| --- | --- | --- | --- | --- | --- | --- |
| Sucrose | | 26.67 | 13.34 | 6.66 | 3.33 | 53.34 |
| AA | M | 0.67 | 0.32 | 0.16 | 0.08 | 1.34 |
|  | H | 0.67 | 0.32 | 0.16 | 0.08 | 1.34 |
|  | V | 0.67 | 0.32 | 0.16 | 0.08 | 1.34 |
|  | R | 0.67 | 0.32 | 0.16 | 0.08 | 1.34 |
|  | T | 0.67 | 0.32 | 0.16 | 0.08 | 1.34 |
|  | W | 0.67 | 0.32 | 0.16 | 0.08 | 1.34 |
|  | I | 0.67 | 0.32 | 0.16 | 0.08 | 1.34 |
|  | L | 0.67 | 0.32 | 0.16 | 0.08 | 1.34 |
|  | F | 0.67 | 0.32 | 0.16 | 0.08 | 1.34 |
|  | K | 0.67 | 0.32 | 0.16 | 0.08 | 1.34 |
| Cholesterol | | 0.17 | 0.17 | 0.17 | 0.17 | 0.17 |
| Lecithin | | 0.17 | 0.17 | 0.17 | 0.17 | 0.17 |
| Ethanol (mL) | | 5 | 5 | 5 | 5 | 5 |
| Vanderzant Vitamin Mix | | 0.5 | 0.5 | 0.5 | 0.5 | 0.5 |
| Wesson Salt Mix | | 0.5 | 0.5 | 0.5 | 0.5 | 0.5 |
| Ascorbic acid | | 1 | 1 | 1 | 1 | 1 |
| Inositol | | 0.25 | 0.25 | 0.25 | 0.25 | 0.25 |
| Choline chloride | | 0.17 | 0.17 | 0.17 | 0.17 | 0.17 |
| Agar | | 3.33 | 3.33 | 3.33 | 3.33 | 3.33 |
| Distilled water (mL) | | 500 | 500 | 500 | 500 | 500 |

**Table S3. The number of spores per unit of surface.** We used a surface regression approach to estimate a parametric nonlinear response surface. This comprises linear ([Carbohydrates] and [Amino acids]) and quadratic components ([Carbohydrates]^2 and [Amino acids]^2) for amino acids and carbohydrate concentration and the cross-product of amino acid and carbohydrate concentration ([Amino acids]*[Carbohydrates]). For the purpose of the analysis, we have scaled all predictors by subtracting their mean value and dividing them by the standard deviation. We considered four different models that were different parameter configurations of the complete model and analyzed their relative performance using AIC-based model selection.

|  | **Number of spores (vol = 0.1µL)** | | | | |
| --- | --- | --- | --- | --- | --- |
| *Predictors* | *Estimates* | *CI* | *Statistic* | *p* | *df* |
| Intercept | 0.01 | -1.16 – 0.11 | 0.09 | 0.925 | 278 |
| [Amino acids] | -0.87 | -1.16 – -0.57 | -5.80 | **<0.001** | 278 |
| [Carbohydrates] | 1.15 | 0.76 – 1.55 | 5.73 | **<0.001** | 278 |
| [Carbohydrates]^2 | -1.64 | -2.15 – -1.12 | -6.27 | **<0.001** | 278 |
| [Amino acids]*[Carbohydrates] | 0.94 | 0.58 – 1.30 | 5.10 | **<0.001** | 278 |
| Observations | 283 | | | | |
| F = 15.59 | P**<0.001** | | | | |
| R^2^ / R^2^ adjusted | 0.183 / 0.171 | | | | |

**Table S4. Fungal surface after 14 days.** We used a surface regression approach to estimate a parametric nonlinear response surface. This comprises linear ([Carbohydrates] and [Amino acids]) and quadratic components ([Carbohydrates]^2 and [Amino acids]^2) for amino acids and carbohydrate concentration and the cross-product of amino acid and carbohydrate concentration ([Amino acids]*[Carbohydrates]). For the purpose of the analysis, we have scaled all predictors by subtracting their mean value and dividing them by the standard deviation. We considered four different models that were different parameter configurations of the complete model and analyzed their relative performance using AIC-based model selection.

|  | **Fungal surface after 14 days** | | | | |
| --- | --- | --- | --- | --- | --- |
| *Predictors* | *Estimates* | *CI* | *Statistic* | *p* | *df* |
| Intercept | 0.01 | -0.07 – -0.09 | 0.27 | 0.790 | 252 |
| [Amino acids] | -1.12 | -1.33 – -0.91 | -10.63 | **<0.001** | 252 |
| [Carbohydrates] | 2.62 | 2.33 – 2.90 | 18.32 | **<0.001** | 252 |
| [Carbohydrates]^2 | -5.17 | -5.68 – -4.65 | -19.76 | **<0.001** | 252 |
| [Amino acids]*[Carbohydrates] | 1.85 | 1.61 – 2.10 | 15.11 | **<0.001** | 252 |
| Observations | 257 | | | | |
| F = 104.6 | P**<0.001** | | | | |
| R^2^ / R^2^ adjusted | 0.624 / 0.618 | | | | |

**Table S5. Fungal surface after 30 days.** We used a surface regression approach to estimate a parametric nonlinear response surface. This comprises linear ([Carbohydrates] and [Amino acids]) and quadratic components ([Carbohydrates]^2 and [Amino acids]^2) for amino acids and carbohydrate concentration and the cross-product of amino acid and carbohydrate concentration ([Amino acids]*[Carbohydrates]). For the purpose of the analysis, we have scaled all predictors by subtracting their mean value and dividing them by the standard deviation. We considered four different models that were different parameter configurations of the complete model and analyzed their relative performance using AIC-based model selection.

|  | **Fungal surface after 30 days** | | | | |
| --- | --- | --- | --- | --- | --- |
| *Predictors* | *Estimates* | *CI* | *Statistic* | *p* | *df* |
| Intercept | 0.00 | -0.06 – -0.07 | 0.15 | 0.882 | 252 |
| [Amino acids] | -1388.70 | -1964.50 – -812.91 | -4.75 | **<0.001** | 252 |
| [Amino acids]^2 | 518.45 | 299.90 – 725.04 | 4.75 | **<0.001** | 252 |
| [Carbohydrates] | 601.95 | 348.72 – 841.26 | 4.76 | **<0.001** | 252 |
| [Carbohydrates]^2 | -1454.03 | -2032.20 – -842.28 | -4.76 | **<0.001** | 252 |
| [Amino acids]*[Carbohydrates] | 809.18 | 468.75 – 1130.97 | 4.76 | **<0.001** | 252 |
| Observations | 258 | | | | |
| F = 146.2 | P**<0.001** | | | | |
| R^2^ / R^2^ adjusted | 0.744 / 0.739 | | | | |

**Table S6. Total number of spores estimated.** We used a surface regression approach to estimate a parametric nonlinear response surface. This comprises linear ([Carbohydrates] and [Amino acids]) and quadratic components ([Carbohydrates]^2 and [Amino acids]^2) for amino acids and carbohydrate concentration and the cross-product of amino acid and carbohydrate concentration ([Amino acids]*[Carbohydrates]). For the purpose of the analysis, we have scaled all predictors by subtracting their mean value and dividing them by the standard deviation. We considered four different models that were different parameter configurations of the complete model and analyzed their relative performance using AIC-based model selection.

|  | **Total number of spores estimated** | | | | |
| --- | --- | --- | --- | --- | --- |
| *Predictors* | *Estimates* | *CI* | *Statistic* | *p* | *df* |
| Intercept | 0.44 | 0.28 – 0.60 | 5.45 | **<0.001** | 2532 |
| [Amino acids] | 0.02 | 0.00 – 0.03 | 2.52 | **0.014** | 253 |
| [Carbohydrates] | 0.00 | -0.01 – 0.01 | 0.22 | 0.829 | 253 |
| [Carbohydrates]^2 | -0.00 | -0.00 – -0.00 | -7.36 | **<0.001** | 253 |
| [Amino acids]*[Carbohydrates] | 0.00 | 0.00 – 0.00 | 6.14 | **<0.001** | 253 |
| Observations | 258 | | | | |
| F = 17.84 | P**<0.001** | | | | |
| R^2^ / R^2^ adjusted | 0.220 / 0.208 | | | | |

**Table S7. Effect of past diet.** We used a linear model to compare the number of secondary spores harvested (per 0.1µL) on a standard medium as a function of the diet used to produce the primary spores (1:199, 1:4 or 2:1).

|  | **Number of spores (vol = 0.1µl)** | |
| --- | --- | --- |
| *Predictors* | *Estimates* | *p* |
| Diet [1:199] vs [2:1] | 1.07 | 0.139 |
| Diet [1:199] vs [1:4] | -0.41 | 0.697 |
| Diet [1:4] vs [2:1] | 1.48 | **0.028** |
| Observations | 16 | |
| F=4.56 | **P=0.03** | |
| R^2^ / R^2^ adjusted | 0.413 / 0.322 | |

|  | **Number of spores (vol = 0.1µL)** | | | | |
| --- | --- | --- | --- | --- | --- |
| *Predictors* | *Estimates* | *CI* | *Statistic* | *p* | *df* |
| Intercept | 0.02 | -0.16 – 0.20 | 0.20 | 0.840 | 122 |
| Diet pairing | 0.05 | -0.02 – 0.13 | 1.42 | 0.157 | 122 |
| Diet | 0.07 | -0.10 – 0.25 | 0.83 | 0.406 | 122 |
| Diet pairing * Diet | 0.02 | -0.05 – 0.10 | 0.60 | 0.547 | 122 |
| Observations | 126 | | | | |
| F=0.92 | P=0.429 | | | | |
| R^2^ / R^2^ adjusted | 0.022 / -0.002 | | | | |

**Table S8. Number of spores in the diet pairing experiment.** Linear model testing the relationship between the number of spores, the diet pairing and the diets. Since the diet pairing * diet was not significant (df = 122, p = 0.57), we reran the model without the interaction to obtain better estimates for the main effects of diet and treatment.

**Table S9. Fungal surface after 14 days in the diet pairing experiment.** Linear model testing the relationship between the area, diet pairings (1:199 vs 2:1, 1:49 vs 2.1, 1:24 vs 2:1, 1:16 vs 2:1, 1:8 vs 2:1) and the diets (1:199 and 2:1). Since the diet pairing * diet was not significant (df = 94, p = 0.63), we reran the model without the interaction to obtain better estimates for the main effects of diet and treatment.

|  | **Fungal surface after 14 days** | | | | |
| --- | --- | --- | --- | --- | --- |
| *Predictors* | *Estimates* | *CI* | *Statistic* | *p* | *df* |
| Intercept | -0.02 | -0.22 – 0.18 | -0.20 | 0.845 | 94 |
| Diet pairing | -0.06 | -0.15 – 0.03 | -1.39 | 0.169 | 94 |
| Diet | -0.07 | -0.27 – 0.14 | -0.65 | 0.519 | 94 |
| Diet pairing * Diet | 0.02 | -0.07 – 0.11 | 0.48 | 0.631 | 94 |
| Observations | 98 | | | | |
| F=0.89 | P=0.448 | | | | |
| R^2^ / R^2^ adjusted | 0.028 / 0.003 | | | | |

**Table S10. Fungal surface after 30 days in the diet pairing experiment.** Linear model testing the relationship between the area, the diet pairing (1:199 vs 2:1, 1:49 vs 2.1, 1:24 vs 2:1, 1:16 vs 2:1, 1:8 vs 2:1) and the diets (1:199 and 2:1). Since the diet pairing * diet was not significant (df = 122, p = 0.35), we reran the model without the interaction to obtain better estimates for the main effects of diet and treatment.

|  | **Fungal surface after 30 days** | | | | |
| --- | --- | --- | --- | --- | --- |
| *Predictors* | *Estimates* | *CI* | *Statistic* | *p* | *df* |
| Intercept | 0.01 | -0.16 – 0.18 | 0.13 | 0.894 | 122 |
| Diet pairing | 0.04 | -0.03 – 0.11 | 1.03 | 0.305 | 122 |
| Diet | -0.35 | -0.51 – -0.18 | -4.12 | **<0.001** | 122 |
| Diet pairing* Diet | 0.03 | -0.04 – 0.10 | 0.893 | 0.354 | 122 |
| Observations | 126 | | | | |
| F=6.68 | **P=0.0003** | | | | |
| R^2^ / R^2^ adjusted | 0.141 / 0.120 | | | | |

**Table S11.** **Survival of ants (infected and uninfected) on different diets.** Cox regression was used to test the effect of the diet and treatment on the survival of the host ants. Survival of the ants was monitored over 64 days. Since the interaction diet * treatment was not significant (LR 2.65, df = 1, p = 0.10), we reran the model without the interaction to obtain better estimates for the main effects of diet and treatment.

|  | **Survival on different diets** | | |
| --- | --- | --- | --- |
| *Predictors* | *LR Chisq* | *df* | *Pr(>Chisq)* |
| Diet | 6.408 | 1 | **0.01** |
| Treatment | 36.180 | 1 | **0.001** |
| Diet * Treatment | 2.65 | 1 | 0.10 |

| Likelihood ratio test = 46.16 | **P=0.0001** |
| --- | --- |

**Table S12. Survival of ants on different diets.** Pairwise comparison with Tukey HSD test: Survival of ants (infected and uninfected) on different diets.

|  | **Survival on different diets** | | | |
| --- | --- | --- | --- | --- |
|  | *Estimates* | *Std. Error* | *z value* | *Pr(>\|z\|)* |
| 1:199 uninfected - 1:199 infected | -1.2235 | 0.2340 | -5.230 | **< 0.001** |
| 1:4 infected - 1:199 infected | 0.1723 | 0.2078 | 0.829 | 0.840 |
| 1:4 uninfected - 1:199 infected  1:4 infected - 1:199 uninfected  1:4 uninfected - 1:199 uninfected  1:4 uninfected - 1:4 infected | -0.5404  1.3958  0.6831  -0.7126 | 0.2118  0.2358  0.2370  0.2101 | -2.551  5.920    2.883  -3.392 | **0.052**  **< 0.001**  **0.020**  **0.004** |

**Table S13. Total Flow (ant per hour) during the collective choice experiment.** To assess the difference in general foraging activity (number of ants foraging), we used a general linear mixed model. The model was fitted by specifying the fixed effects: treatment, and week and the random effects (colony). Following the Shapiro-Wilk test, we obtained the following values: *w* = 0.91 and *p* <0.001 implying that the distribution of the data was significantly different from normal distribution. Total flow was then normalized using the “*bestNormalize*” function (“*bestNormalize*” package).

|  | **Total Flow (ant per hour)** | | |
| --- | --- | --- | --- |
| *Predictors* | *Estimates* | *CI* | *p* |
| Treatment | -0.04 | -0.18 – -0.11 | 0.615 |
| Week | -0.43 | -0.53 – -0.33 | **<0.001** |
| Treatment * Week | 0.14 | 0.04 – 0.24 | **0.005** |
| **Random Effects** | | | |
| σ^2^ | 0.57 | | |
| τ_00_ _Colony_ | 0.01 | | |
| ICC | 0.01 | | |
| N _Colony_ | 22 | | |
| Observations | 110 | | |
| Marginal R^2^ / Conditional R^2^ | 0.429 / 0.436 | | |

**Table S14. Flow of ants per minute on each branch during the collective choice experiment.** To assess the difference in the ants’ distribution between the two diets, we used generalized linear mixed models. The model was fitted by specifying the fixed effects: treatment and the random effect (colony) and the error family (binomial).

|  | **Flow of ants per minute on each branch** | | |
| --- | --- | --- | --- |
| *Predictors* | *Odds Ratios* | *CI* | *p* |
| Treatment | 1.71 | 1.49 – 1.96 | **<0.001** |
| **Random Effects** | | | |
| σ^2^ | 3.29 | | |
| τ_00_ _Colony_ | 0110 | | |
| ICC | 0.03 | | |
| N _Colony_ | 22 | | |
| Observations | 110 | | |
| Marginal R^2^ / Conditional R^2^ | 0.078 / 0.108 | | |

**Table S15. Ratio of amino acids and carbohydrate collected during the collective choice experiment.** To assess the difference in the ratio of amino acids to carbohydrate collected per individual ant between the two treatments (infected *vs* uninfected), we used a linear mixed model. The model was fitted by specifying the fixed effects: treatment and the random effect (colony).

|  | **Ratio of amino acids to carbohydrate collected** | | |
| --- | --- | --- | --- |
| *Predictors* | *Estimates* | *CI* | *p* |
| Treatment | -10.72 | -17.63 – -3.81 | **0.002** |
| **Random Effects** | | | |
| σ^2^ | 339.00 | | |
| τ_00_ _ColonyID_ | 0.00 | | |
| N _ColonyID_ | 22 | | |
| Observations | 110 | | |
| Marginal R^2^ / Conditional R^2^ | 0.078 / NA | | |

**Table S16. Ant mortality during the collective choice experiment.** To assess the difference in the number of dead ants per week between the two treatments (infected *vs* uninfected), we used a linear mixed model. The model was fitted by specifying the fixed effects: treatment, week and the random effect (colony).

|  | **Mortality (number of dead ants per week)** | | |
| --- | --- | --- | --- |
| *Predictors* | *Estimates* | *CI* | *p* |
| Treatment | 18.42 | 6.24 – 30.60 | **0.003** |
| Week | -2.13 | -4.59 – 0.33 | 0.089 |
| Treatment * Week | -1.59 | -4.92 – 1.73 | 0.347 |
| **Random Effects** | | | |
| σ^2^ | 157.09 | | |
| τ_00_ _ColonyID_ | 37.74 | | |
| ICC | 0.19 | | |
| N _ColonyID_ | 22 | | |
| Observations | 110 | | |
| Marginal R^2^ / Conditional R^2^ | 0.253 / 0.398 | | |

**Table S17**. Immune gene expression. Candidate gene approach was applied to test if the β−1,3-glucan injection elicits an immune response in ants. The expression level of 4 insect immune genes was measured by droplet digital PCR (ddPCR). Expression levels of the immune gene were normalized to the housekeeping gene (*GAPDH*) before further statistical analysis.

|  | **Immune genes**  ***Hymenoptaecin*** | |
| --- | --- | --- |
| *Predictors* | *Estimates* | *p* |
| (Intercept) | 0.22 | 0.213 |
| Treatment | -1.05 | **0.001** |
| Group | 0.50 | **0.013** |
| Observations | 68 | |
| R^2^ / R^2^ adjusted | 0.367 / 0.348 | |

**Table S18**. Immune gene expression. Candidate gene approach was applied to test if the β−1,3-glucan injection elicits an immune response in ants. The expression level of 4 insect immune genes was measured by droplet digital PCR (ddPCR). Expression levels of the immune gene were normalized to the housekeeping gene (*GAPDH*) before further statistical analysis.

|  | **Immune genes**  ***Serpin 27a* (SPN-27a)** | |
| --- | --- | --- |
| *Predictors* | *Estimates* | *p* |
| (Intercept) | 0.04 | 0.83 |
| Treatment | 0.51 | 0.03 |
| Group | 0.36 | 0.12 |
| Observations | 68 | |
| R^2^ / R^2^ adjusted | 0.10 / 0.08 | |

**Table S19**. Immune gene expression. Candidate gene approach was applied to test if the β−1,3-glucan injection elicits an immune response in ants. The expression level of 4 insect immune genes was measured by droplet digital PCR (ddPCR). Expression levels of the immune gene were normalized to the housekeeping gene (*GAPDH*) before further statistical analysis.

|  | **Immune genes**  **Defensin** | |
| --- | --- | --- |
| *Predictors* | *Estimates* | *p* |
| (Intercept) | -0.13 | 0.55 |
| Treatment | 0.21 | 0.38 |
| Group | 0.05 | 0.82 |
| Observations | 68 | |
| R^2^ / R^2^ adjusted | 0.01 / -0.01 | |

**Table S20**. Immune gene expression. Candidate gene approach was applied to test if the β−1,3-glucan injection elicits an immune response in ants. The expression level of 4 insect immune genes was measured by droplet digital PCR (ddPCR). Expression levels of the immune gene were normalized to the housekeeping gene (*GAPDH*) before further statistical analysis.

|  | **Immune genes**  **Relish** | |
| --- | --- | --- |
| *Predictors* | *Estimates* | *p* |
| (Intercept) | 0.24 | 0.25 |
| Treatment | -0.61 | 0.01 |
| Group | 0.08 | 0.72 |
| Observations | 68 | |
| R^2^ / R^2^ adjusted | 0.09 / 0.07 | |

**Table S21**. Individual choice assay of (*i*) Triton-X treated, (*ii*) *Metarhizium*-infected, (*iii*) microinjected with physiological saline solution (i*v*) microinjected with β−1,3-glucan ants.

|  | | *Chisq* | *Pr (>Chisq)* |
| --- | --- | --- | --- |
| *Metarhizium* infected vs Zymosan injected ants | | 0.49 | 0.48 |
| *Metarhizium* infected vs Triton ants | | 4.15 | 0.04 |
| Zymosan injected vs Saline treated ants | | 6.67 | 0.009 |

**Table S22**. Primer sequences used for droplet digital PCR (ddPCR), amplification lengths and annealing temperatures (*Defensin 1, Relish, HYM = Hymenoptaecin* and *SPN* = *Serpin 27a*, *GAPDH* = housekeeping gene).

| **Genes** | **Sequence** | **Amplicon length** | **Annealing temperature** |
| --- | --- | --- | --- |
| Defensin 1 | F: 5’-GCTGATTTACGTTTCAGCCG  P:5’[FAM]CGGAGCCGGCGCGTGACCTG[BHQ1]  R: 5’-ATTGCCAGGACAGAAGATCG | 152 | 60°C |
| HYM | F: 5’-GTTTTCAACAATGGCCGCAC  P:5’[FAM]CGCACGTCGGCCTTCAGGCGG[BHQ1]  R: 5’-GAAACTGGGTTTGACCTCTG | 142 | 60°C |
| SPN | F: 5’-GGCTTCCATACGATGGTTCA  P:5’[FAM]TGGCGAGAGACATATGGGCCATGCAAGAGT[BHQ1]  R: 5’-GGGATCCAAACATCGAGAGG | 151 | 60°C |
| Relish | F: 5’-ACCATACATCAGCGTCTTGG  P:5’[FAM]TGGGACCTTGGTTGGAAGTGGACGCAG[BHQ1]  R: 5’-CGTATTATCGCGTTCTCCGT | 165 | 60°C |
| GAPDH | F: 5’-AGATTGCTGTCTTCAGCGAG  P:5’[HEX]ACAGGCGTGTTCACCACCATCGATAAAGCC[BHQ1]  R: 5’-AATGACTTTCTTCGCACCGC | 143 | 60°C |

**Supplementary figures**

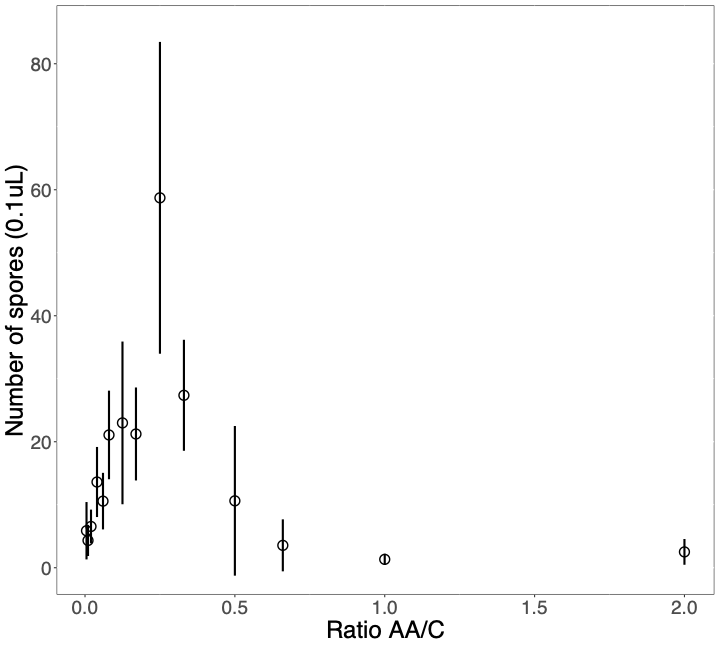

**Figure S1**. **No-choice diet experiment (relating to Figure 1A).** The effect of ratio AA:C on spore production. The number of spores per µL (volume = 0.1µL) was determined for fungi confined for 30 days to 1 of 18 diets varying in the ratio of amino acids to carbohydrates.

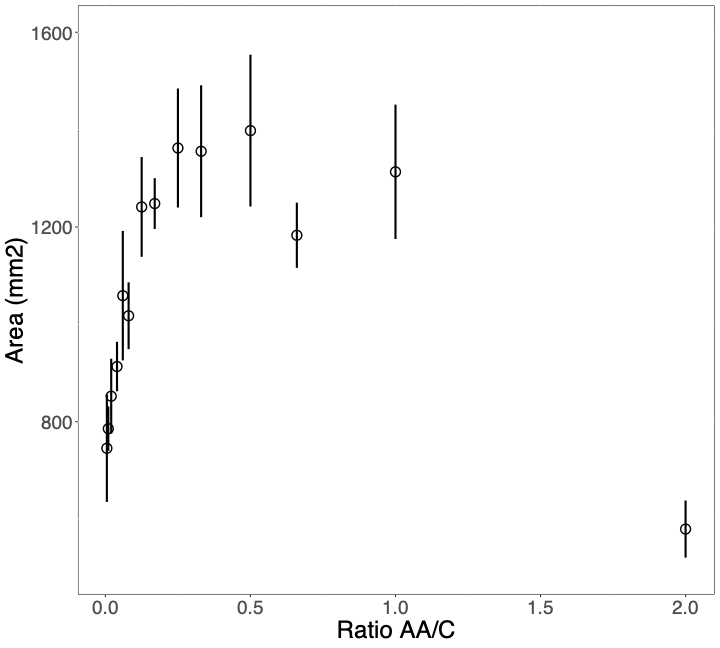

**Figure S2. No-choice diet experiment (relating to Figure 1C).** The effect of ratio AA:C on fungal growth (area: day 14). Fungal surfaces (mm^2^) were quantified for fungi confined for 14 days to 1 of 18 diets varying in the ratio of amino acids to carbohydrates.

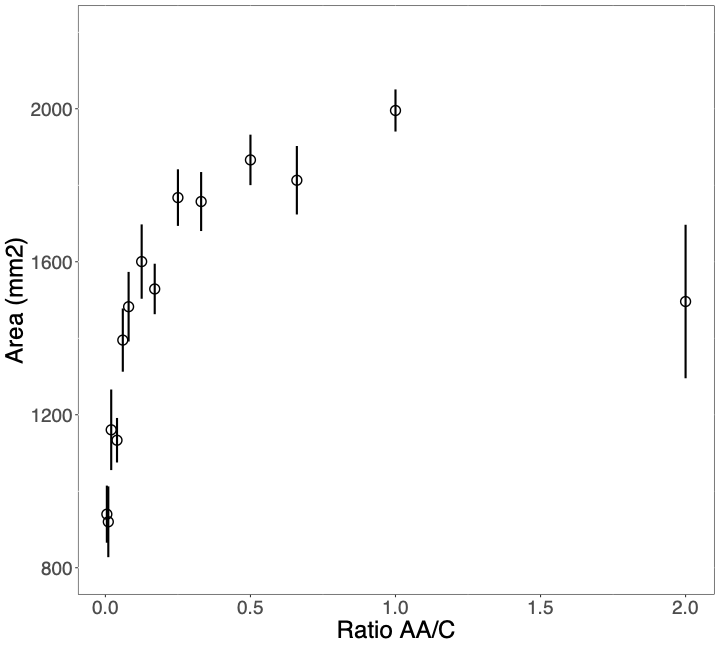

**Figure S3. No choice diet experiment (relating to Figure 1D).** The effect of ratio AA:C on fungal growth (area: day 30). Fungal surfaces (mm^2^) were quantified for fungi confined for 30 days to 1 of 18 diets varying in the ratio of amino acids to carbohydrates.

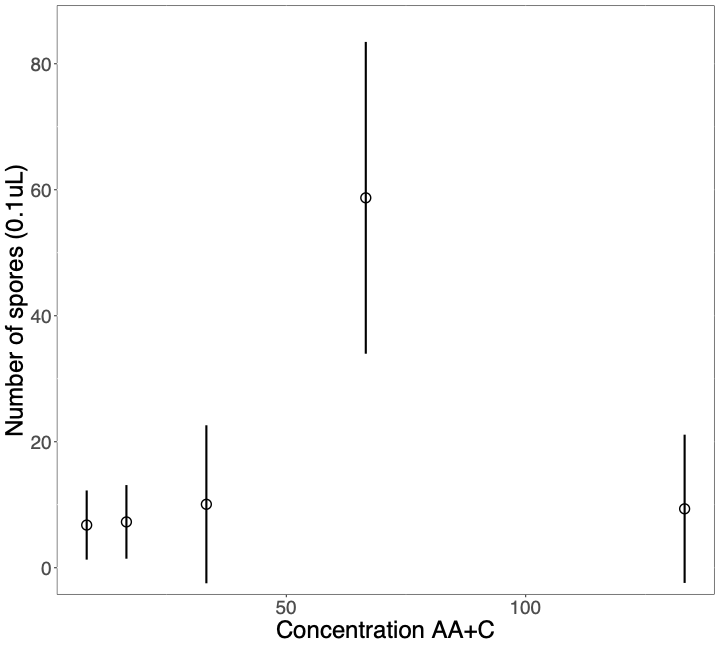

**Figure S4. Dilution experiment (relating to Figure 1A).** The effect of total AA+C concentration on spore production. The number of spores per µL (volume = 0.1µL) was determined for fungi confined for 30 days to 1 of 5 diets varying in the concentration of amino acids to carbohydrates. The following diets were used: control: undiluted 1:4, diluted 1:4/2, 1:4/4, 1:4/8, concentrated 1:4/x2.

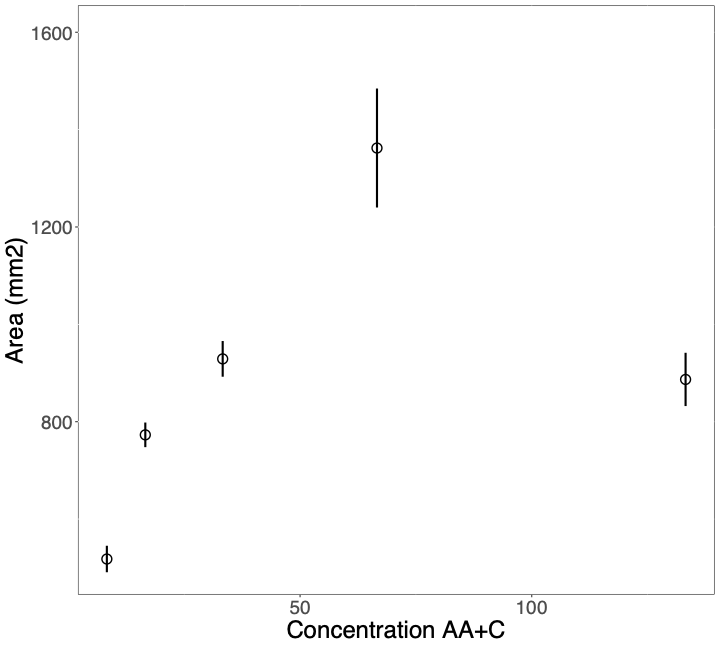

**Figure S5. Dilution experiment (relating to Figure 1B).** The effect of total AA+C concentration on fungal growth (area: day 14). *M. brunneum* growth (mm^2^) was quantified for fungi confined for 14 days to 1 of 5 diets varying in the ratio of amino acids to carbohydrates. The following diets were used: control: undiluted 1:4, diluted 1:4/2, 1:4/4, 1:4/8, concentrated 1:4/x2.

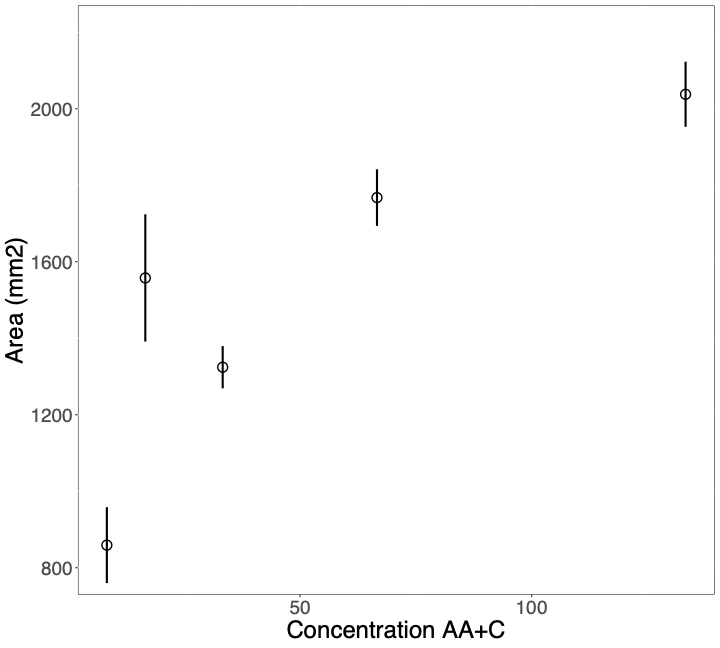

**Figure S6. Dilution experiment (relating to Figure 1C).** The effect of total AA+C concentration on fungal growth (area: day 30). *M. brunneum* growth (mm^2^) was quantified for replicates confined for 30 days to 1 of 5 diets varying in the ratio of amino acids to carbohydrates. The following diets were used: control: undiluted 1:4, diluted 1:4/2, 1:4/4, 1:4/8, concentrated 1:4/x2.

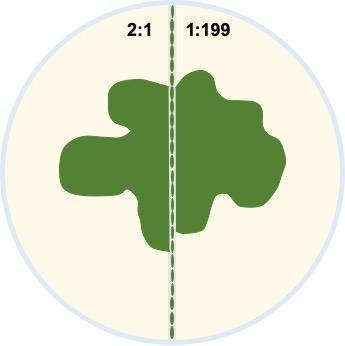

**Figure S7. Schematic representation of the fungal diet choice experiment.** The Petri dishes (diameter 55 mm) were divided into two parts, one of the diets 2:1 was poured on one side of the Petri dish (5 mL). The second diet 1:199 was poured on the other side of the Petri dish (5 mL). To each Petri dish, 20 µl of a 2 x 10^7^ spores ml^−1^ suspension of *Metarhizium brunneum* was applied in a line (dark green), in the centre of a Petri dish directly in contact with both diets.
